# Cold atmospheric plasma for improved sustainability and fish welfare in recirculating aquaculture systems

**DOI:** 10.64898/2026.11.23.768699

**Authors:** Martina Balazinski, Veronika Hahn, Christin Höhne, Robert Wagner, Anja Poehlein, Gerd-Michael Arndt, Klaus-Dieter Weltmann, Mirko Basen, Juergen F. Kolb

**Affiliations:** Leibniz Institute for Plasma Science and Technology, Felix-Hausdorff Str. 2, 17489 Greifswald; Mecklenburg-Vorpommern Research Centre for Agriculture and Fisheries (LFA MV), Institute of Fisheries, Research Station Aquaculture, Südstr. 8, Born 18375, Germany; Genomic and Applied Microbiology & Göttingen Genomics Laboratory, Georg-August-University, Göttingen, Germany; Microbiology, Institute of Biological Sciences, Faculty of Mathematics and Natural Sciences, University of Rostock, Albert-Einstein-Str. 3, 18059 Rostock, Germany; Department of Maritime Systems, Interdisciplinary Faculty, University of Rostock, Albert-Einstein-Str. 21, 18059 Rostock, Germany; Interdisciplinary Faculty of the University of Rostock, Institute of Physics, Albert-Einstein-Str. 23-24, 18059 Rostock, Germany

## Abstract

Decontamination of water is mandatory for the cultivation of aquatic species in recirculating aquaculture systems (RAS). Avoiding economic and ecological losses while ensuring safe conditions is important for the future introduction of new technologies. Therefore, the implementation of cold atmospheric plasma (CAP) within RAS was examined for the first time.

For the integration of this technology two equal RAS were set up, one using CAP, the other one ultraviolet light (UV) for water decontamination. Experiments with Rainbow trout (*Oncorhynchus mykiss*) were performed for three weeks to compare CAP with UV as reference. The CAP device was switched on intermittently for 8 hours daily (with intervals of 10 min on and 30 min off), while two UV lamps were turned on continuously throughout the entire time of 3 weeks, which required more energy. Fish development was tracked by fish weight and stress levels (cortisol measurements). Values for pH, conductivity, oxygen, temperature, microbial load, microbiome and ion composition were also monitored.

Microbial loads for both setups were reduced by four orders of magnitude. The biodiversity of the microbiome of the two technologies was similar with a ß-diversity between 19 and 25. Deviations within each setup were low with the α-diversity ranging from 2.23-2.93 (CAP) and 2.42-2.79 (UV). The water monitoring revealed similar results for pH value, conductivity, oxygen and temperature. No differences for the cortisol measurements and also the fish weight were determined.

The results demonstrate CAP as a competitive and more energy efficient alternative to UV benefiting fish welfare and development in a similar way.

The equal development of fish and water parameters and low energy consumption compared to UV makes CAP an energy efficient tool, likewise enhancing fish welfare.

## 1. Introduction

Fish and fish products provide an important contribution to a healthy diet. The consumption of fish is expected to further increase in the following years (21.1 kg in 2022-2024 to 21.8 kg in 2034 per capita; OECD 2025). The demand can no longer be met by fishing alone and for the alternative production in aquacultures annual growth rates between 4.5% and 7.3% are expected till 2032 (Global Market Insights 2024) and 2034 (OECD 2025). The OECD predicts a total share of aquacultures of 56% by then (OECD 2025). Especially the aquaculture production by recirculating aquaculture systems (RAS) offers the potential to mitigate challenges of open cultures affected by climate changes, environmental constraints and impacts and strain on natural resources. In addition, easier to control aspects of biosecurity water-quality management and the opportunity for continuous, off-season production become strategic values (Hashmi et al. 2025).

However, special attention must be paid to a healthy cultivation and decontamination technologies applied that, prevent the outbreak of diseases by the inactivation of fish-pathogenic microorganisms in the water (Li et al. 2023, Aly and Fathi 2024). In this context also antibiotic-resistant microorganisms (ARM) become an increasing issue of increasing importance (Teitge et al. 2020). Decontamination needs have to be balanced against constraints imposed by fish welfare, which is ideally not impaired. The load as well as the diversity of the microorganisms is of relevance, as too high numbers pose the risk of fish-pathogenic microorganisms to thrive. Fluctuations in the microbiome might promote imbalances favouring microbial strains which cause stress or diseases (Xiong et al. 2019). Accordingly, a balanced, intact microbiome plays an essential role to ensure a healthy development of the fish, necessary for a safe and nutritious product (Infante-Villamil et al. 2019, Roques and van de Vis 2025).

Although, UV and ozone have been effectively applied as decontamination technologies in aquaculture systems (Sharrer and Summerfelt 2007) drawbacks, such as high energy costs and increasing microbial resistances require the development and integration of new technologies as addition or alternative (Badiola et al. 2012). Moreover, UV does not penetrate far into the water, especially turbid water, limiting the antimicrobial efficacy (Gullian et al. 2012). Conversely, ozone, as a strong oxidizing agent can pose risks to animals and humans (Penru et al. 2013).

In these regards, cold atmospheric plasma (CAP) can serve as alternative, which has already been successfully demonstrated for the inactivation of fish-pathogenic microorganisms, relevant in RAS (Balazinski et al. 2025). Its antimicrobial efficacy, especially against harmful Gram-negative microorganisms has been shown to be comparable to that of UV. The technology is based on physico-chemical interactions, the generation of reactive oxygen and nitrogen species (RONS) in water, offering the opportunity to provide sufficient decontamination independent from water turbidity. The life-time of the reactive species is rather short ensuring a safe application in contrast to ozone.

The safety was further determined *in vitro* by toxicity and mutagenicity studies (Balazinski et al 2026). These studies revealed safe boundary conditions for the tested fish cell line (RTgill-W1) as well as no mutagenicity for the performed Ames test. However, the gained data referred to laboratory tests performed *in vitro*, thus lacking necessary information for living fish.

For this reason, CAP was now implemented in the real environment of RAS and evaluated as a decontamination technology. The pilot investigation was setup with two equally equipped systems. The CAP device was integrated in one setup, while UV lamps were used in the control setup for decontamination purposes. For the cultivated stock, Rainbow trout (*Oncorhynchus mykiss*) served as a species common in RAS production.

The main concern of this study was dedicated to fish welfare. Regardless, the control of the microbial load played a significant role. Concurrently, changes to the microbiome were evaluated. In addition, the stress hormone cortisol, water parameters, fish behaviour weight development of the fish provided a comprehensive set of parameters to assess impact on fish welfare. Thus, the approach and scope of this study directly supports and addresses policy recommendations, that were elaborated by the European Commission to improve the cultivation strategies for the five most farmed fish (Atlantic salmon, Rainbow trout, Common carp, Gilthead seabream and European sea bass) in the future (European Commission 2021, Pavlidis et al. 2023).

## 2. Materials and methods

### 2.1 Experimental setup for CAP and UV treatment

The technology of CAP as a decontamination technology in RAS was compared to UV as a common standard technology.

#### 2.1.1 RAS

Two setups with two fish tanks each, equipped with drum and biofilter differed solely by the decontamination method, which was either CAP or UV (Fig. 1). The water exchange was adjusted equally in both systems accordingly to the naturally increasing nitrate values due to rising stocking densities.

**Fig. 1.**
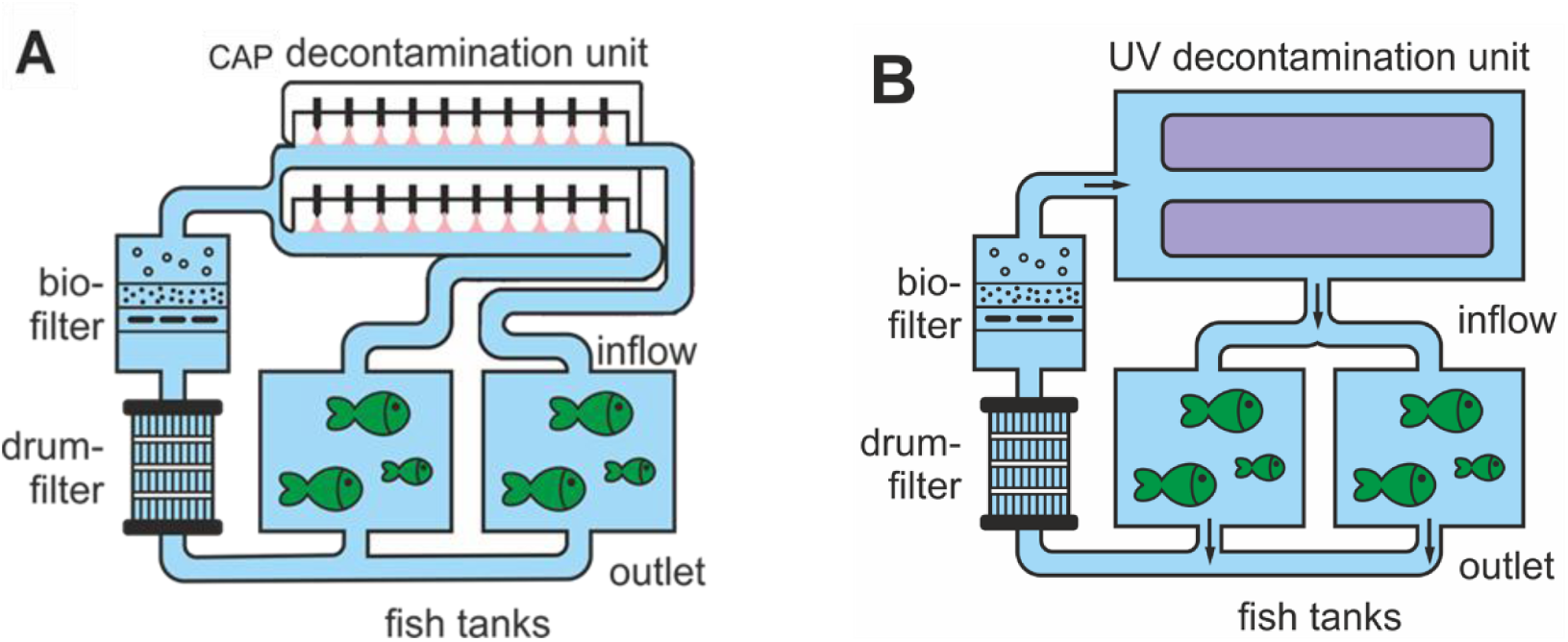
RAS of (A) CAP or (B) UV as decontamination method.

In each fish tank of 90l a stock of roughly 960 g Rainbow trout was placed, resulting in approximately 74 fish per tank, with an average initial weight of 13 g. The initial stock density was roughly 10 kg/m³. In each setup, the water was pumped from the tanks through a drum filter passing the biofilter, before reaching the decontamination unit. After flowing through the decontamination unit the water returned back to the tanks. The flow rate for both setups was 270 l/h.

The duration of the experiment was set to three weeks, which was supposed to result in the doubling of the fish weight. The fish were transferred to the tanks one week prior to the beginning of the experiment to allow for an adaption of the fish to the system. Until the start of the experiment the decontamination in both setups was ensured by UV treatment.

#### 2.1.2 Decontamination units

The CAP decontamination unit consisted of two parallel acrylic pipes (Fig. S I, Supplementary Information). The water was run through these pipes, simultaneously being treated by CAP, provided by electrode pins placed above the water surface. The ignition of CAP was achieved by 10 transformers, each applying an AC modulated (50 Hz) high voltage of 24 kV (peak-to-peak) to a pair of sharpened stainless-steel electrodes. A constant distance between the electrode and the water surface ensured a pin-to-liquid discharge with consistent treatment (Fig. 1A). The CAP device was operated for 10 min, followed by a 30 min pause during a total treatment time of 8 h (from 7 am to 3 pm), resulting in a treatment of 2.02 h per day.

The UV decontamination unit consisted of two UV lamps which were arranged in parallel within one pipe, each with a power input of 55 W (Fig. 1B). The water was flushed through the pipe and distributed to the tanks after treatment. For the UV treatment the lamps were as it is common for RAS switched on for 24 h during the entire trial of three weeks.

#### 2.1.3 Sampling

For the determination of the total viable cell count (2.2.1), 16S rRNA gene amplicon sequencing (2.2.2) the ion composition (2.3.1) and the cortisol levels (2.4.3), water samples were taken at different points of the RAS setups (Fig. 2).

**Fig. 2.**
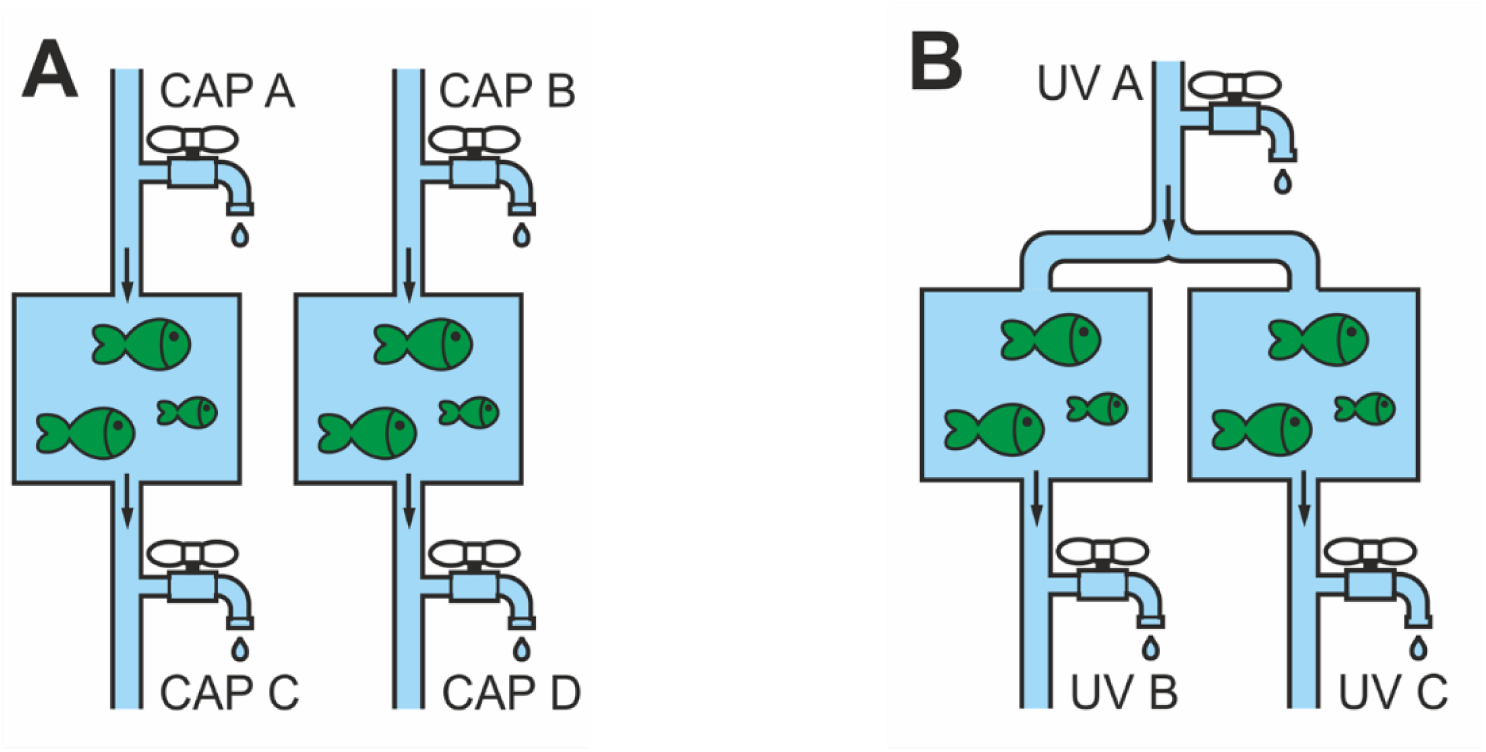
Section of RAS with a detailed view on the sampling points of (A) the CAP setup with inflow (CAP A and CAP B) and outlet (CAP C and CAP D) as well as for (B) the UV setup with inflow (UV A) and outlet (UV B and UV C).

Therefore, the inflow samples (CAP A and B; UV A) were taken right after the respective decontamination unit, before the water enters the tank. The outflow samples (CAP C and D; UV B and C) allowed for sampling without disturbing the fish, which could affect cortisol levels.

Samples for the different analytics, i.e., total viable count 2.2.1, 16S rRNA gene amplicon sequencing 2.2.2, ion composition 2.3.1, cortisol levels 2.4.3) various sampling days and sampling times were used for the CAP and UV setup (Tab. 1).

**Tab. 1.** Sampling days and sampling times (t_0_: prior to 7 am, t_1_: after 7 am, t_2_: after 3 pm) for both setups for different analytical methods, i.e., TVC (total viable count), IC (ion-exchange chromatography), M (16S rRNA gene amplicon sequencing, C (cortisol levels).

| **Day of sampling** | **t<sub>0</sub>** | **t<sub>1</sub>** | **t<sub>2</sub>** |
|---|---|---|---|
| **0** | TVC, IC, M, C | C | TVC, IC |
| **2** | TVC |  |  |
| **7** | TVC, IC, M, C | C | TVC, IC |
| **9** | TVC |  |  |
| **14** | TVC, IC, M, C | C | TVC, IC |
| **16** | TVC |  |  |
| **21** | TVC, IC, M, C |  |  |

### 2.2 Microbiological examinations

The evaluation of the microbial load (2.2.1) as well as the determination of species via MALDI-TOF (2.2.2) and 16S rRNA gene amplicon sequencing (2.2.3) were conducted for both setups CAP and UV to compare the decontamination properties and likewise the impact on the microbial community.

#### 2.2.1 Determination of the total viable cell count

For the determination of the microbial load in the water samples, 1 mL from the respective sampling points (Fig. 2) was spread undiluted on agar plates containing tryptic soy agar (TSA, Carl Roth, Karlsruhe, Germany). The inoculated plates were incubated at 30°C for up to 5 days. The viable cell count was determined by manual colony counting (colony-forming units, cfu/ml).

The viable count was analysed twice a week (Tab. 1), with sampling dates at the beginning of the experiment, right before the CAP treatment was started (time point t_0_, prior to 7 am) and at the end of the CAP treatment (time point t_2_, after 3 pm) at the same day. Another sampling was done two days later (at time point t_0_) for additional interim results in between of the major sampling dates each week. Samples of the UV setup were taken accordingly, serving as control and reference.

Final samples were taken at the end of the trial on day 21.

#### 2.2.2 16S rRNA gene amplicon sequencing

For the determination of the microbiome in both settings (CAP and UV) water samples were taken at the beginning of the experimental trial (2.1.3, Tab. 1), followed by weekly samplings. Therefore, 2 l of the water from each RAS were taken from each respective inflow and outflow point. The samples were centrifuged, the supernatant discarded and the remaining pellet was resuspended in 2 ml 0.9% sodium chloride solution (Carl Roth, Karlsruhe, Germany) and stored at -20°C until further processing.

The respective samples were extracted using the DNeasy® PowerSoil® Pro Kit (Qiagen, Venlo, The Netherlands). All procedures were performed in accordance to the manufacturer’s guideline (Qiagen 2023). After the extraction the quantity and quality of the DNA was determined by a spectrophotometer (Implen NanoPhotometer, Munich, Germany) by measuring the absorbance at 260 nm, 280 nm and 230 nm.

Bacterial communities were assessed by amplification and sequencing of the V3-V4 region of the 16S rRNA gene. In addition to the respective samples, a positive control (from *Escherichia coli*), a negative control (nuclease-free water (Invitrogen, Thermo Fisher Scientific, Dreieich, Germany)) and a mock sample (ZymoBIOMICS Microbial Community Standard, Zymo Research, Orange, USA) as quality control standard were included. For the polymerase chain reaction (PCR) the following parameters were applied according to a protocol by Engelhardt et al. (Böer et al. 2024): Initial denaturation at 95°C for 1 min, followed by 25 cycles of 98°C for 45 s, 55°C for 45 s and 72°C for 30 s, concluding with 72°C for 5 min, before cooling down to 10°C. Following the amplification, the DNA was purified by using the GeneRead Size Selection Kit (Qiagen, Venlo, The Netherlands) according to the manufacturer’s guideline (Qiagen 2020). The 16S rRNA gene amplification was evaluated by gel electrophoresis for correct amplicon size of 550 base pairs (bp). In addition, the PCR products were quantified and a total volume of 40 µl, with a minimum concentration of 25 ng/µl was used for Illumina sequencing. Library preparation, sequencing and analysis were performed as described by Böer et al., with the exception that the SILVA SSU database v138.2 has been used for taxonomic assignment.

Operational taxonomic units (OTU) and amplicon sequencing variants (ASV) from the Illumina sequencing were queried against the SILVA database using the Basic Local Alignment Search Tool (BLAST) heuristic. The results of the BLAST were stratified by excluding low abundances (relative abundances lower than 2%). The relative abundances used in this study were recalculated based on the stratified data.

#### 2.2.3 MALDI-TOF

Colonies from the water samples of both setups (CAP and UV), which grew on TSA agar plates were analysed by MALDI-TOF by the company LABOKLIN (Bad Kissingen, Germany). Therefore, colonies of different morphology, like colour, shape and size, were selected from all samples taken during the experiment.

#### 2.2.4 Biodiversity calculations

Biodiversity in the RAS water was assessed by calculating the Shannon index *H*′ (Pielou 1966) for the α-diversity and the β-diversity according to Whittaker (Whittaker 1960) based on the relative abundances of each determined species from the 16S rRNA amplicon sequencing.

The Shannon index for n species found in one site is defined as:

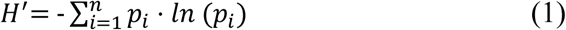

With *p_i_* quantifying the relative abundance of species i.

The Whittaker β-diversity between two sites (in this case the CAP-vs. the UV-treated tank) is defined as the degree of change in community composition. Similar to (Baselga 2010) and (Legendre 2007), the change in community composition can be expressed as:

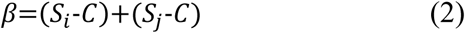

With *S_i_* and *S_j_* quantifying the number of different genera found in site I and j, respectively, and C the number of common species (intersection) in site i and j.

*S_i_* = Number of different genera found in site i

*S_j_* = Number of different genera found in site j

*C* = Intersection of found species in site i and j

### 2.3 Analyses of water parameters

The microbial testing was complemented by an analysis of ions and chemical compounds, such as nitrate, and of pertinent water parameters, such as pH value, conductivity and oxygen concentration to monitor changes and ensure safe conditions throughout the experimental trial.

#### 2.3.1 Ion composition

The anionic components were analysed using an ion-exchange chromatography Dionex ICS 6000 (Thermo Fisher Scientific, Dreieich, Germany). The anions chloride, carbonate, sulphate, nitrate and nitrite were detected via conductivity. For the anion separation a Dionex IonPac AS18 column with a pre-connected Dionex IonPac AG18 guard column (both Thermo Fisher, Dreieich, Germany) was used. The eluent was derived from ultrapure water produced from a GenPure Pro Barnstead unit (Thermo Fisher Scientific, Dreieich, Germany) in combination with a potassium hydrogen solution cartridge, providing a 23 mM concentrated solution. Each sample was measured twice, resulting in two technical replicates.

#### 2.3.2 Conductivity, pH value, oxygen and temperature

For the inline monitoring the water parameters pH value, oxygen concentration and conductivity were measured and logged every 15 min with a data logger (Hach Lange GmbH, Berlin, Germany). In addition, the respective values were recorded manually each day for the sampling times t_0_ and t_2_.

### 2.4 Fish-welfare monitoring

The assessment of the fish welfare was based on observations of fish behaviour, the development of the fish weight, the stress levels expressed by the cortisol concentrations found in water samples and also by the changes of the microbiome.

#### 2.4.1 Fish behavior

The behaviour of the fish was regularly observed by experienced staff to ensure fish welfare and evaluated according to scoresheet ratings for the gill movement, swimming activity and the overall behaviour. These provided also instructions of action (see Tab. S I, Supplementary Information).

#### 2.4.2 Fish weight

For the fish development the weight of 20 randomly selected fish per tank (for both setups) was noted each week. The fish were weighed right after the water samples were taken for the microbiological examinations and the cortisol concentration measurements.

#### 2.4.3 Cortisol levels

The stress level was determined by weekly measurements of the stress hormone cortisol found in water samples. Therefore, water was taken before and after starting the CAP treatment (t_0_ and t_1_). Sampling was performed approximately 5 min before the CAP treatment was started and right after it ended. The sample preparation was performed accordingly to (Martinez 2014). Therefore, remaining particles were removed by a glass filter of 1 l of water sample from the tank outlet. Afterwards the sample was extracted by an Oasis water HLB Plus Short Cartridge (Waters, Milford, USA). The cartridge had to be activated with 5 ml of methanol (Carl Roth, Karlsruhe, Germany) prior to use. After the extraction of the 1 l sample the cartridges were cleaned with 5 ml of distilled water. The cartridges were then sealed and stored in a freezer at -20°C. For the cortisol concentration measurements, the extract was eluted with 5 ml ethyl acetate (Carl Roth, Karlsruhe, Germany) from the thawed cartridge and evaporated under nitrogen gas atmosphere. The gained extract was taken up in 0.5 ml phosphate buffered saline mixed with ethanol (5%) and bovine albumin serum (0.1%; all Carl Roth, Karlsruhe, Germany), which was adjusted to a pH value of 7.2.

The actual cortisol measurement was carried out according to the instruction of the manufacturer (IBL International GmbH 2024) for the Cortisol Saliva ELISA Testkit (IBL International GmbH, Hamburg, Germany). The cortisol content was determined at a wavelength of 450 nm with a TECAN Infinite 200 Pro M Nano^+^ spectrophotometer (Tecan Group AG, Männedorf, Switzerland). Measurements were carried out with two technical replicates.

### 2.5 Energy consumption

For the comparison of the efficiency of both tested decontamination technologies the energy consumption of the CAP device was determined by connecting the device to a performance measuring device (PCE Instruments, Meschede, Germany). Values in the range of 480-520 W were measured, which refer to the power applied for plasma generation, but also for auxiliary circuitry, such as cooling ventilators.

## 3. Results

### 3.1 Microbial inactivation

The microbial load of the two setups (CAP and UV) was determined for the respective inflow and outlet sampling points and different days (2.1.3, Tab. 1).

Thus, for the evaluation of the mean microbial load per setup the average of the inflow and outlet was considered. In general, the calculated average microbial load of the samples taken over three weeks showed similar results for both setups (Fig. 3). The average load (throughout the entire duration of the trial) for the CAP setup resulted in 2.16·10^4^ cfu/ml, while the value of the UV setup was slightly higher with 3.27·10^4^ cfu/ml.

**Fig. 3.**
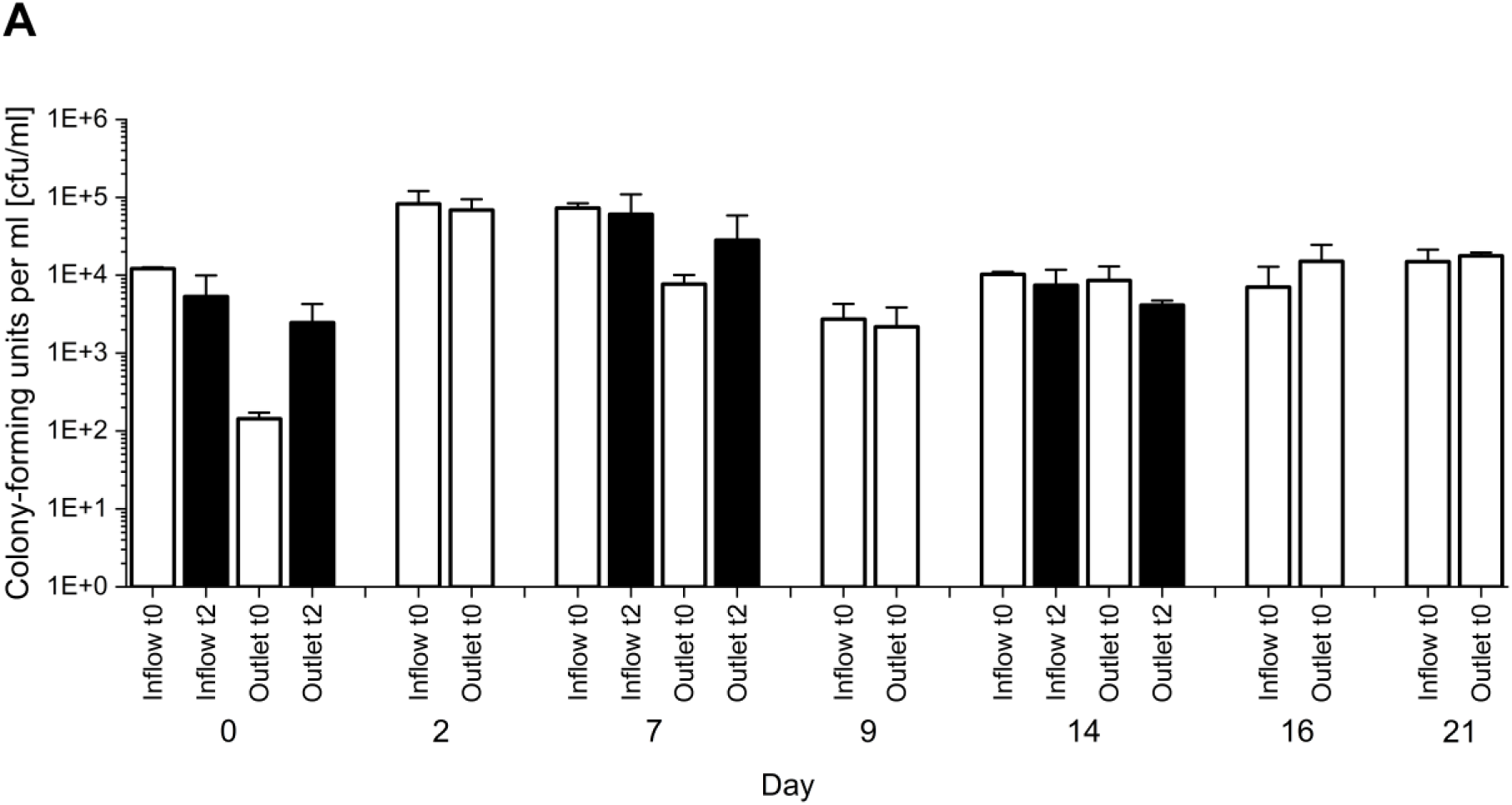
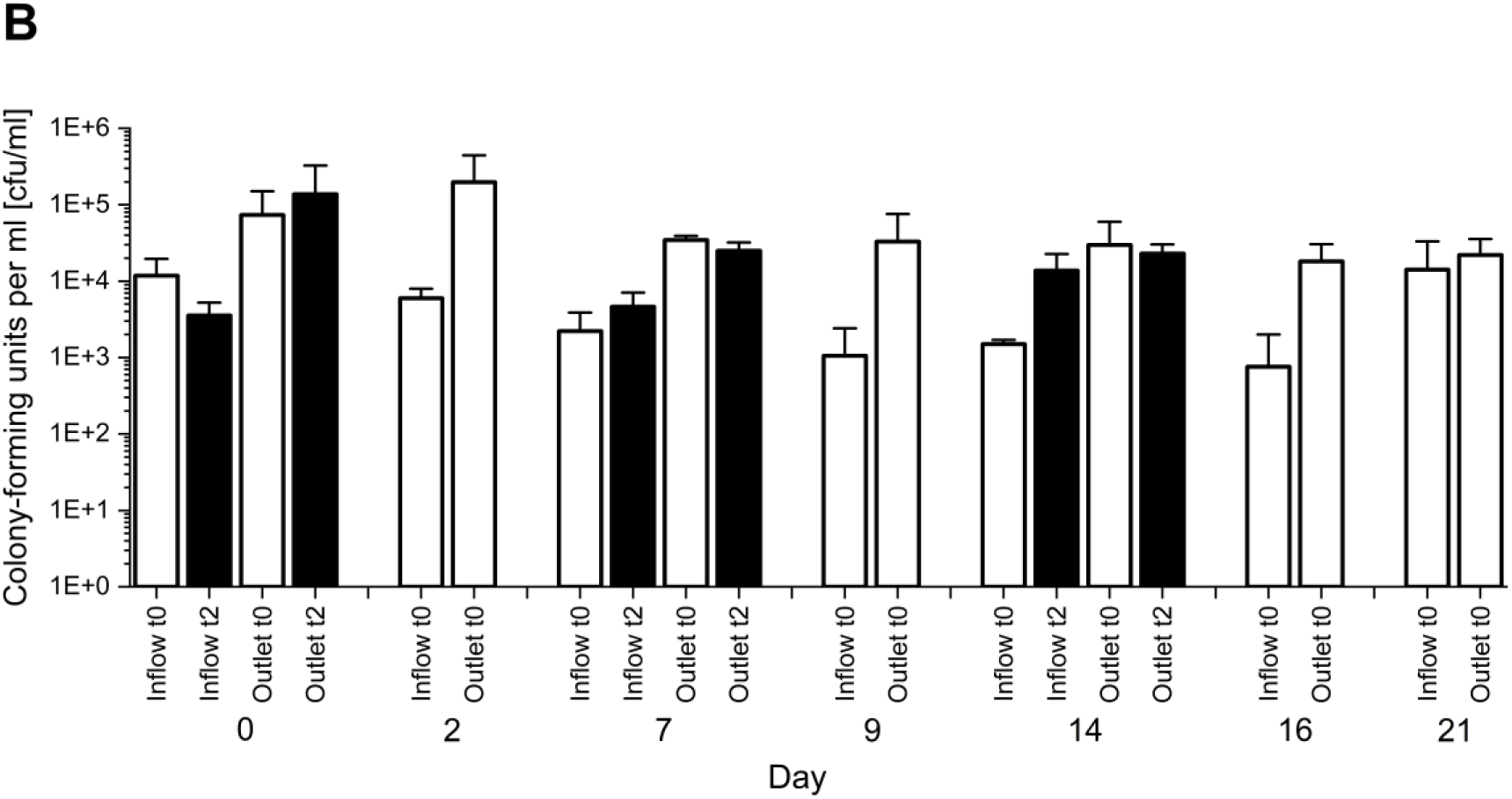
Averages and standard deviations for 2 technical replicates of the microbial load (as cfu/ml) of the water samples from (A) the CAP setup and (B) the UV setup at the inflows (average of CAP A and B, UV A) and the outlets (average of CAP C and D, average of UV B and C) for the sampling times t_0_ (white columns) and t_2_ (black columns) of the respective sampling days.

Prior to the start of the trial the decontamination in both setups was ensured by UV treatment for one week, necessary for the fish to adapt. Therefore, the initial microbial loads are reflected by the samples taken on day 0 for t_0_. Subsequently, for the CAP setup (Fig. 3A) differences in the viable cell count were detected for the different sampling days and sampling times (t_0_ and t_2_). While at the beginning of the experiments (day 0) the average microbial load from the inflow and outlet position was 5.02·10^3^ cfu/ml, an increase of almost one order of magnitude up to 5.35·10^4^ cfu/ml was detected for the sampling dates on day 2 and 7. The following samples (from day 9 on) whereas showed a lower microbial load (9.77·10^3^ cfu/ml). The comparison of the different sampling positions (inflow and outlet) and sampling times shows, that except for the results of day 0, the differences in the microbial load were similar. Additionally, the results gained for the sampling times t_0_ and t_2_, representing the sampling before the CAP device was switched on and after an eight hours treatment period, showed also no remarkable differences.

For the detected microbial loads (from day 0 to 21) of all sampling points (inlet and outflow) only little deviations of less than one order of magnitude (0.65) resulted for the mean.

For the decontamination via UV, consistent microbial loads would be expected within a trial, as it was already applied in the adaption phase of the fish. Interestingly, the microbial loads detected at the first sampling dates (day 0 and 2) differed between the sampling points. Thus, the inflow samples had a load of 7.12·10^3^ cfu/ml, the outlet samples resulted in a roughly one order of magnitude higher value (1.37·10^5^ cfu/ml). In the continuation of the experiment (from day 9 on) the values between the sampling points aligned, resulting in an average load of 1.60·10^4^ cfu/ml. In these regards, automated steering processes regulating fresh water supply must be considered as a factor for fluctuating microbial loads. However, the microbial load detected for the respective sampling points followed a clear pattern. The load at the inflow position was always lower than at the outlet. For the different sampling times (t_0_ and t_2_) only slight deviations were detected. In total the microbial loads were similar in both setups (in particular after day 2).

### 3.2 Microbial community

#### 3.2.1 Microbiome analyses

The microbiome analyses were performed to investigate, whether the CAP treatment affects the microbial community in a different way than UV. Therefore, a comparison of the general phyla and further genera distribution was done. The results were plotted for the respective decontamination unit and consecutive sampling days (see Tab. S II, Supplementary Information). The distribution of the phyla showed, that the abundances of *Pseudomonadota* decreased by 56% in the CAP setup (Fig. 4 top) especially within the first week of the experiment (from 55% to 24%), whereas the phyla *Chloroflexota* and *Verrucomicrobiota* increased during the same period (*Chloroflexota* by 40% and *Verrucomicrobiota* by 85%). In the UV setup (Fig. 4 bottom), the relative abundance of *Pseudomonadota* remained similar for the three weeks of experiment (57% ± 2%), while the abundance of *Bacillota* decreased after one week showing a 3.5 times reduction in week two and three compared to the beginning (8%).

**Fig. 4.**
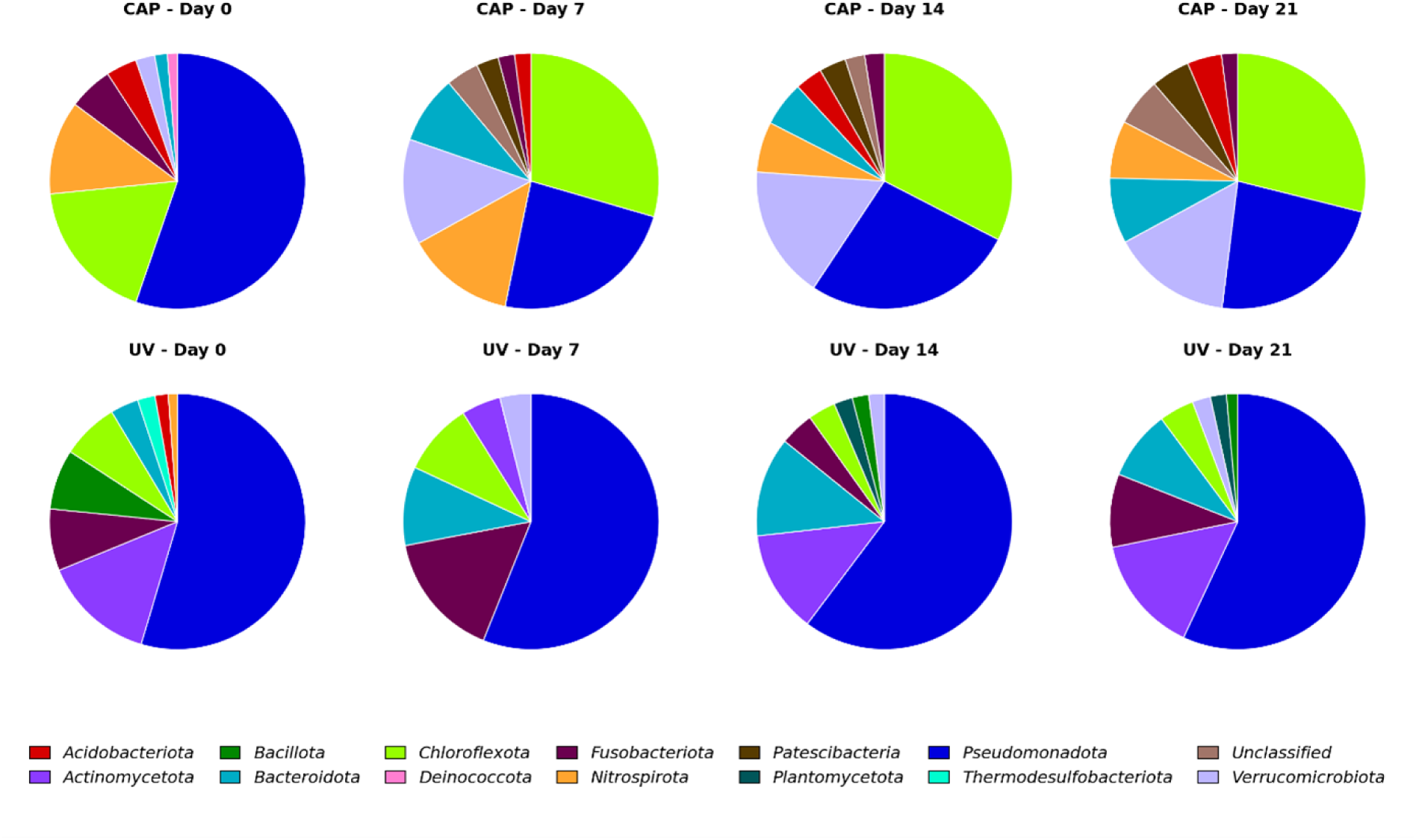
Relative abundances of the detected phyla of the stratified data for CAP (left) and UV (right) of the respective sampling days (see 2.1.3).

For the assessment of the microbiome within the respective setup (CAP or UV) the number of the detected genera in the stratified data served for the evaluation of the biodiversity within the respective treatment applied (Tab. 2).

**Tab. 2.**
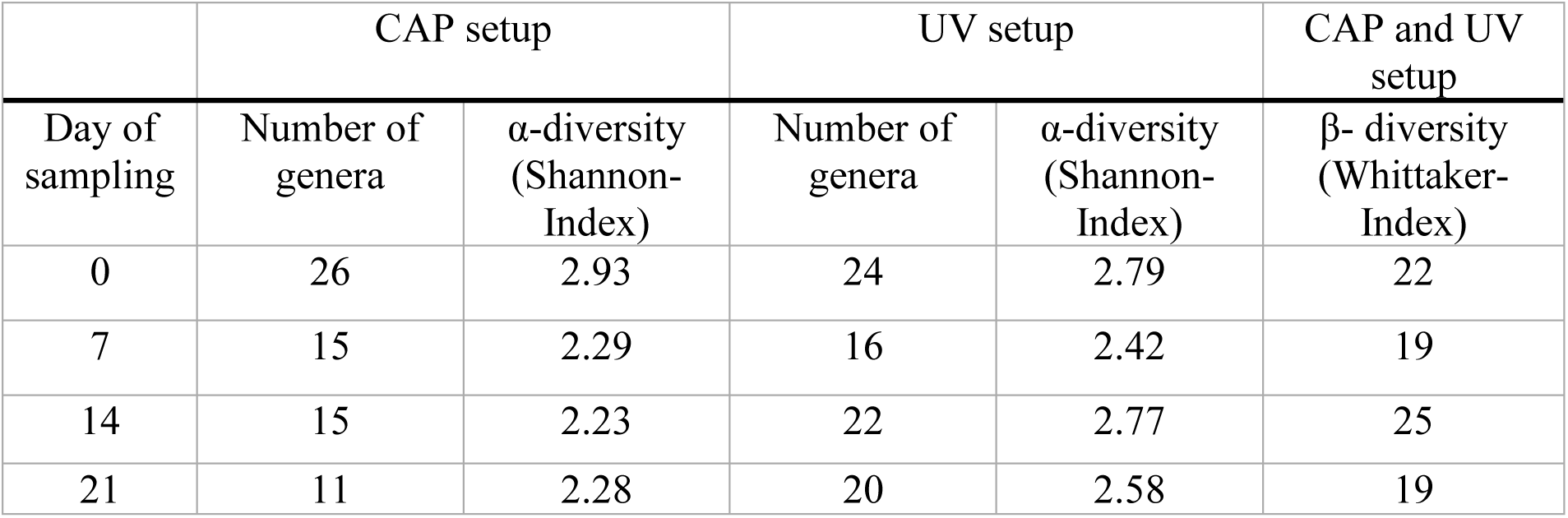
Summation of the detected genera in the stratified data and the calculated values for the α- and ß-diversity per sampling day as the average of the sampling points for the CAP and UV setup.

|  | CAP setup |  | UV setup |  | CAP and UV setup |
|---|---|---|---|---|---|
| Day of sampling | Number of genera | α-diversity (Shannon-Index) | Number of genera | α-diversity (Shannon-Index) | β- diversity (Whittaker-Index) |
| 0 | 26 | 2.93 | 24 | 2.79 | 22 |
| 7 | 15 | 2.29 | 16 | 2.42 | 19 |
| 14 | 15 | 2.23 | 22 | 2.77 | 25 |
| 21 | 11 | 2.28 | 20 | 2.58 | 19 |

At the beginning of the experiment the number of detected genera was slightly different for the two setups with 26 for CAP and 24 for UV. After one week a striking reduction was detected for both setups. For CAP treatment the number of genera was reduced to 15 (by 42%), while for UV the decrease to 16 detected genera (by 35%) was less. In the following weeks these numbers remained stable for the CAP setup, while increasing again to 22 and 20 was determined.

The corresponding changes in the biodiversity within the experiment duration and treatment applied were calculated by the Shannon index (Tab. 2). At the beginning of the experiments the Shannon indices for CAP and UV were nearly the same (CAP: 2.93, UV: 2.79). In the following weeks the Shannon index for the CAP treatment showed reduced and stable values of 2.29, 2.23 and 2.28. For the UV setup, also a reduction was noted to 2.42 within the first week, while the values increased to 2.77 and 2.58 during the following weeks.

In addition to the Shannon index for α-diversity also the Whittaker index, describing β-diversity, was calculated to compare the microbial diversities for both treatments (CAP and UV). The value for the Whittaker index at the beginning of the trial was 22, followed by 19, 25 and 19 in the weeks afterwards. Thus, the values showed no major fluctuations of the diversity between the microbiome of CAP and UV.

The derivation of the indices for α- and β-diversity was complemented by an assessment of the distribution of genera in both systems. The indices provide insight into the impact of the decontamination units on the microbial communities in the water. In general, a multitude of genera was detected indicating a very complex microbial community (Fig. S II, Supplementary Information). However, the uncertainty in the taxonomic classification for the genera named *Incertae* must be considered. Nevertheless, the biggest changes in the microbial community occurred in the first week for both setups. At the beginning, the *Incertae* genus assigned to the phylum *Chloroflexota* together with the genera *Nitrospira* and *Pseudomonas* showed the highest relative abundances (15.4%, 11.9% and 10.0%). By day 7 the relative abundances of *Incertae* assigned to the phylum *Chloroflexota* almost doubled (to 28.8%), while *Nitrospira* slightly increased (to 13.9%). The genera with the next highest abundances were now *Luteolibacter* and *Fuscovulum* (13.5% and 10.2%). For the rest of the trial this community remained similar. In contrast, the microbial community in the UV setup showed a different distribution. Thus, at the beginning the genera *Pseudomonas*, *Rhodococcus* and *Cetobacterium* were dominantly present (21.0%, 10.9% and 7.8%) a decrease of almost 90% was detected for *Rhodococcus* after the first week (1.4%). On the contrary, the unclassified genus assigned to *Comamonadaceae* increased by a factor of 3 (5.4% to 16.1%), *Cetobacterium* doubled (to 15.9%), while *Pseudomonas* slightly decreased (to 18.7%). In the following weeks the unclassified genus remained stable, while the relative abundance of *Pseudomonas* more than halved. Further fluctuations in the genera distribution along the trial occurred.

The comparison of the both setups, revealed that three genera were consistently found at each sampling day: *Cetobacterium*, *Flavobacterium* and *Pseudomonas.* Thus, the abundances of these genera fluctuated within the respective setup, as well as the sampling days (see Fig. 5).

**Fig. 5.**
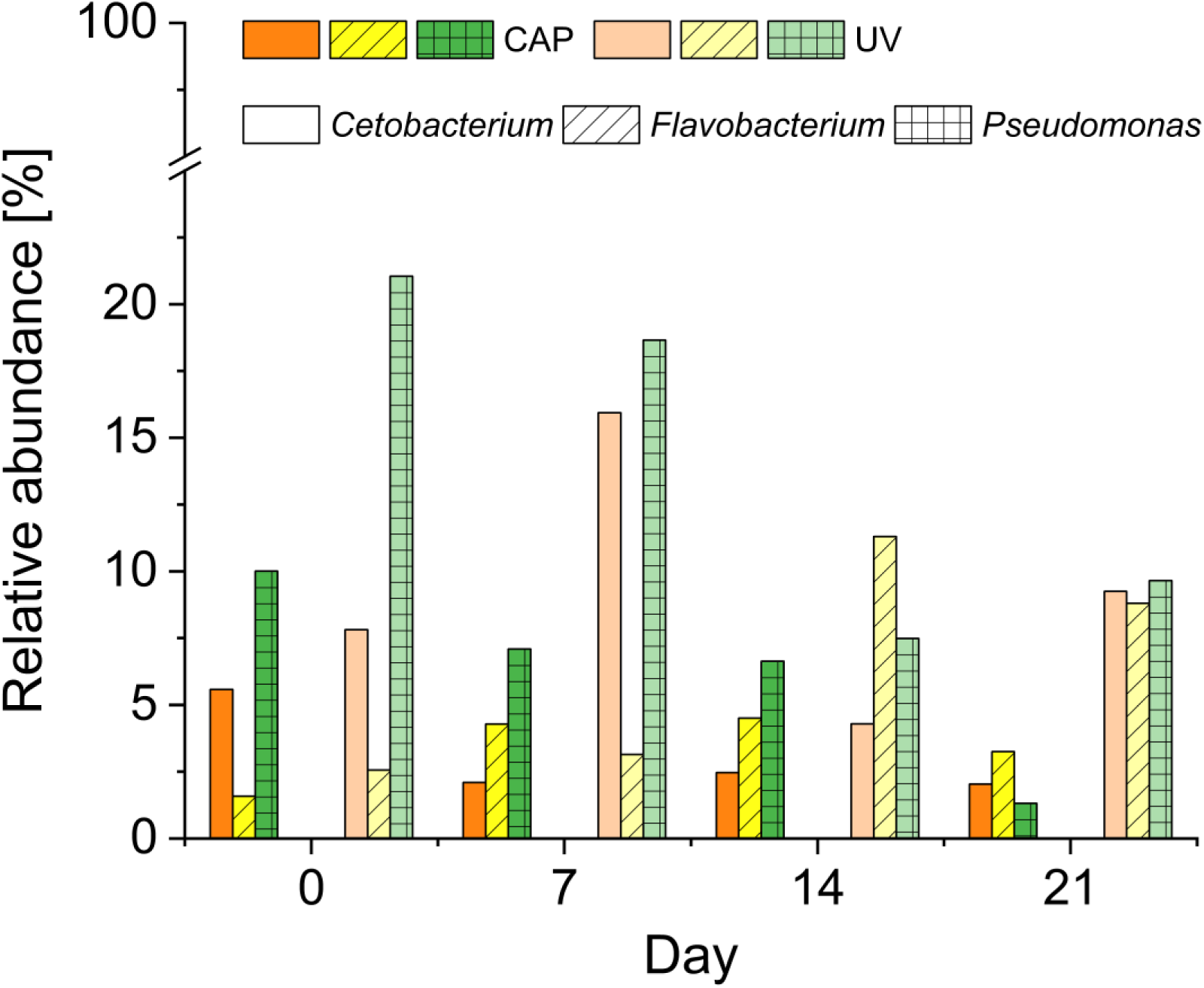
Relative abundances of the detected genera *Cetobacterium, Flavobacterium* and *Pseudomonas* for (A) CAP and (B) UV at the sampling days 0, 7, 14 and 21.

For the CAP setup an increase of the abundance of the genus *Flavobacterium* was detected until day 14 (from 1.6% to 4.5%). For the genera *Cetobacterium* and *Pseudomonas* a decrease throughout the trial was noted. Starting with a relative abundance of 5.6% for *Cetobacterium* by day 0 with a reduction of approximately 64% to 2.0% by day 21. An even higher drop was found for *Pseudomonas* with a decline of even 87% from day 0 with 10.0% to 1.3% by day 21.

For the UV setup, the initial relative abundance of 2.6% for *Flavobacterium* stayed similar by day 7, rising 3.5-times by day 14 (to 11.3%), followed by a reduction of 22% on day 21 (to 8.8%). In comparison, a high initial value for *Pseudomonas* (21.0%) was determined. However, the relative abundance decreased (to 18.7%) on day 7 and even stronger (to 7.5%) on day 14, followed by an increase at day 21 (to 9.7%). The relative abundances of *Cetobacterium* fluctuated during the trial. By day 7 an increase by a factor of 2 (from 7.8% to 15.9%) was detected, which then decreased by a factor of 3.7 on day 14 (to 4.3%). On the last day the relative abundance increased to 9.3% again, resulting in a little higher value compared to day 0.

#### 3.2.2 Species determination

MALDI-TOF served for the identification of microorganisms grown on culture media (TSA plates to isolate and identify individual species) from both setups. Next to the species identification, the microorganisms were taxonomically categorized (Tab. 3). In total thirteen microorganisms were cultivated and isolated on TSA, which were assigned to ten genera, most of them were part of the phylum *Pseudomonadota*, with the genera *Pseudomonas*, *Roseomonas*, *Serratia*, *Paracoccus*, *Acinetobacter* and *Acidovorax*. The genera *Streptococcus* and *Carnobacterium* were representative for the phylum *Bacillota*. The genera *Chryseobacterium* and *Exiguobacterium* were assigned to the phyla *Bacteroidota* and *Bacillota*.

**Tab. 3.** Categorization of microorganisms isolated after CAP and UV treatment by MALDI-TOF analyses

| Phylum | Family | Species |
|---|---|---|
| *Bacillota* | *Bacillaceae* | *Exiguobacterium mexicanum* |
|  | *Carnobacteriaceae* | *Carnobacterium maltaromaticum* |
|  | *Streptococcaceae* | *Streptococcus canis* |
| *Bacteroidota* | *Weeksellaceae* | *Chryseobacterium scophthalmum* |
| *Pseudomonadota* | *Comamonadaceae* | *Acidovorax facilis* |
|  | *Moraxellaceae* | *Acinetobacter johnsonii* |
|  | *Paracoccaceae* | *Paracoccus yeei* |
|  | *Pseudomonadaceae* | *Pseudomonas anguilliseptica* |
|  | *Pseudomonadaceae* | *Pseudomonas corrugata* |
|  | *Pseudomonadaceae* | *Pseudomonas koreensis* |
|  | *Pseudomonadaceae* | *Pseudomonas libanensis* |
|  | *Acetobacteraceae* | *Roseomonas mucosa* |
|  | *Yersiniaceae* | *Serratia grimesii* |

### 3.3 Water condition

Water parameters (ion composition, pH value, conductivity and oxygen) were monitored to ensure fish welfare and also to determine a correlation of changes with microbial inactivation and composition of the microbiome.

#### 3.3.1 Ion composition

Especially anions, such as nitrate and nitrite are important parameters for water quality and were quantified together with chloride, carbonate and sulphate (Fig. 6).

**Fig. 6.**
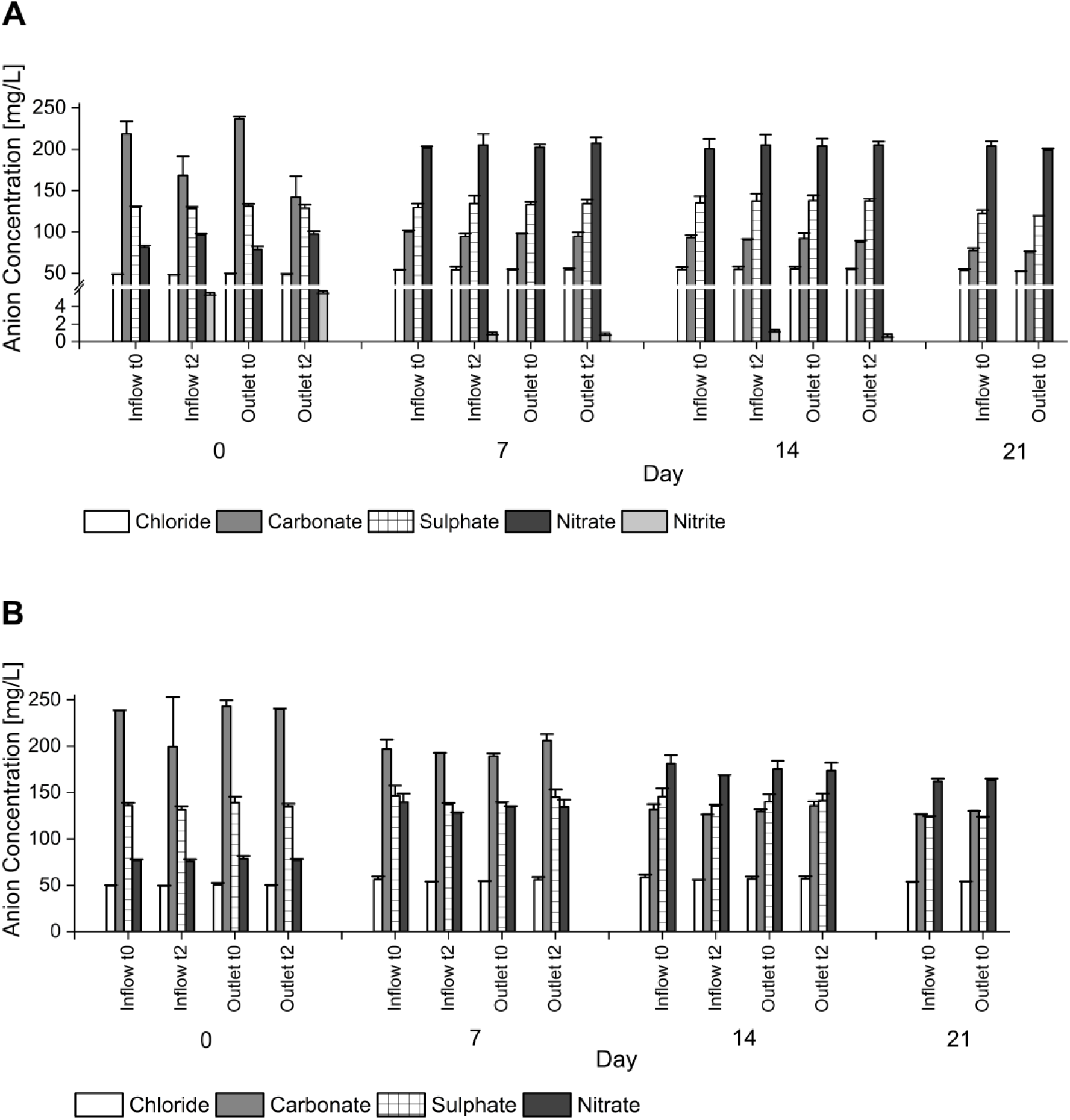
Averages and standard deviations of the anion concentrations (chloride, carbonate, sulphate, nitrate and nitrite) for 2 technical replicates of the water samples from (A) the CAP setup and (B) the UV setup at the inflows (average of CAP A and B, UV A) and the outlets (average of CAP C and D, average of UV B and C) for the sampling times t_0_ and t_2_ of the respective sampling days.

For the CAP setup (Fig. 6A) the concentrations before treatment (at t_0_ of day 0), for chloride were 48.69 mg/l, for carbonate 227.71 mg/l, for sulphate 130.49 mg/l, and 79.76 mg/l for nitrate. No nitrite was detected. After CAP treatment at the sampling time t_2_, carbonate concentrations dropped from 202.11 mg/l by one third (to 155.25 mg/l), while the nitrate concentrations increased by approximately 20% (to 97.02 mg/l). Nitrite concentration had increased to 5.71 mg/l. No relevant changes were observed for concentrations of sulphate and chloride (with an only moderate change for chloride from 48.29 mg/l to 54.09 mg/l after the first week). These concentrations also did not change for consecutive sampling days 7, 14, and 21. Carbonate concentrations stayed after a decrease to 99.13 mg/l within one week, at this level up to the end of the experiment. This was also observed for nitrate but with concentrations increasing 2.5 times to 202.11 mg/l, which were about maintained during the remainder of the experiment. Notably, the measured concentrations were independent from sampling point (inflow, outlet) or sampling time (t_0_, t_1_) after the first week, except for nitrite, which only and consistently showed up with a concentration of approximately 1 mg/l after each treatment of 8 hours, probably due to its utilization as food source by the biofilter.

As expected, for the UV setup, the respective concentrations were similar, i.e., for t_0_ at day 0 (Fig. 6B). Values, similar to the CAP treatment, were also found for the concentrations of sulphate (stayed constant) and chloride (which increased by 10% from 50.14 mg/l to 55.24 mg/l within a week). Interestingly, carbonate concentration decreased by about 20% to 193.10 mg/l in a week and further to 130.85 mg/l, hence, less than for the CAP treatment, until the end of the trial. The most notable differences were observed for concentrations of nitrate, which increased from 77.95 mg/l to 137.21 mg/l by day 7 and to 178. 43 mg/l to the end of the experiment. Accordingly, they were then about 12% lower than for the CAP treatment. However, no nitrite at all was detected for the UV treatment.

#### 3.3.2 Water quality parameters

Standard water parameters, that are relevant to ensure stable cultivation conditions of fish were continuously monitored. These included pH values, dissolved oxygen, temperature and conductivity (Fig. S III, Supplementary Information). No differences were observed between the CAP and UV setup. The pH values started at a high level of almost 9 and decreased within 4 days to a mean of 8. Values for the dissolved oxygen were on average about 10 mg/l in a range between 8 and 12 mg/l. An outlier in the oxygen level was noted on day 8 for the UV setup due to a change of position of the sensor. The conductivity in both systems was between 750 and 850 µS/cm and the temperature ranged from 16 to 20°C.

### 3.4 Fish welfare

#### 3.4.1 Fish behaviour

No abnormal fish behaviour was observed throughout the experiment for both setups (CAP and UV). Especially, no changes in the gill movement, the swim and overall behaviour were detected (Tab. S I, Supplementary Information).

#### 3.4.2 Fish weight

The weight of the fish almost doubled from approximately 18 g up to 35 g during the three weeks of the experiment (Fig. 7), resulting in a rough increase from 10 to 20 kg/m^3^ of the stocking density-a level corresponding to the expected and deliberately chosen stocking density, which is common for raising fish trout in recirculating aquaculture systems. Since, juvenile Rainbow trout was used in our study this development was expected. The increase in mass was similar for CAP and UV treatment.

**Fig. 7.**
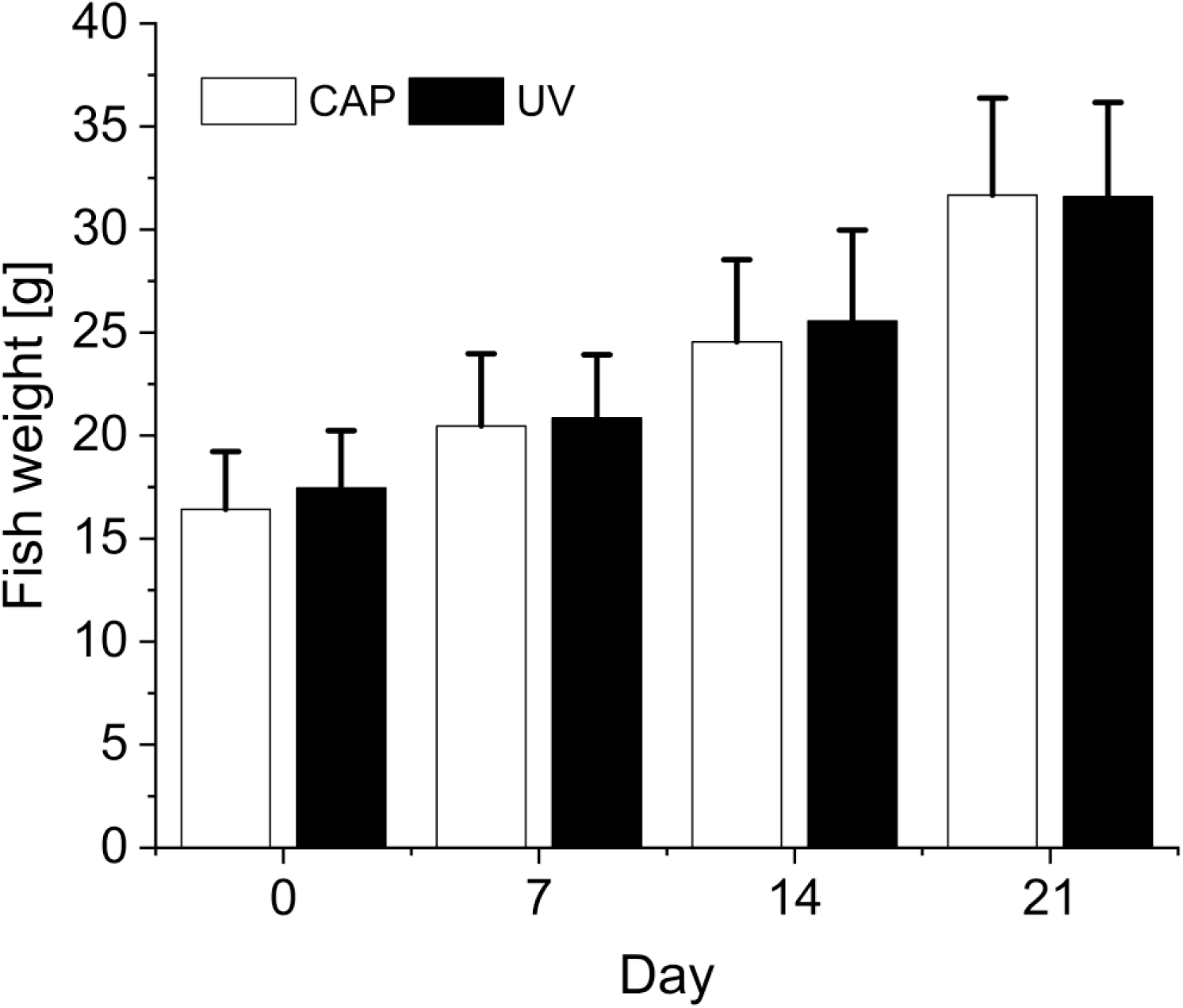
Averages and standard deviations for 20 biological replicates of the fish weight of day 1, 7, 14 and 21 of the experimental trial for the decontamination technologies CAP and UV.

#### 3.4.3 Cortisol level

The impact of the CAP and UV operation on fish was analysed by cortisol measurements in the water at the respective outlet sampling points. The cortisol concentration was measured right before switching on the CAP device and then immediately afterwards (Fig. 8).

**Fig. 8.**
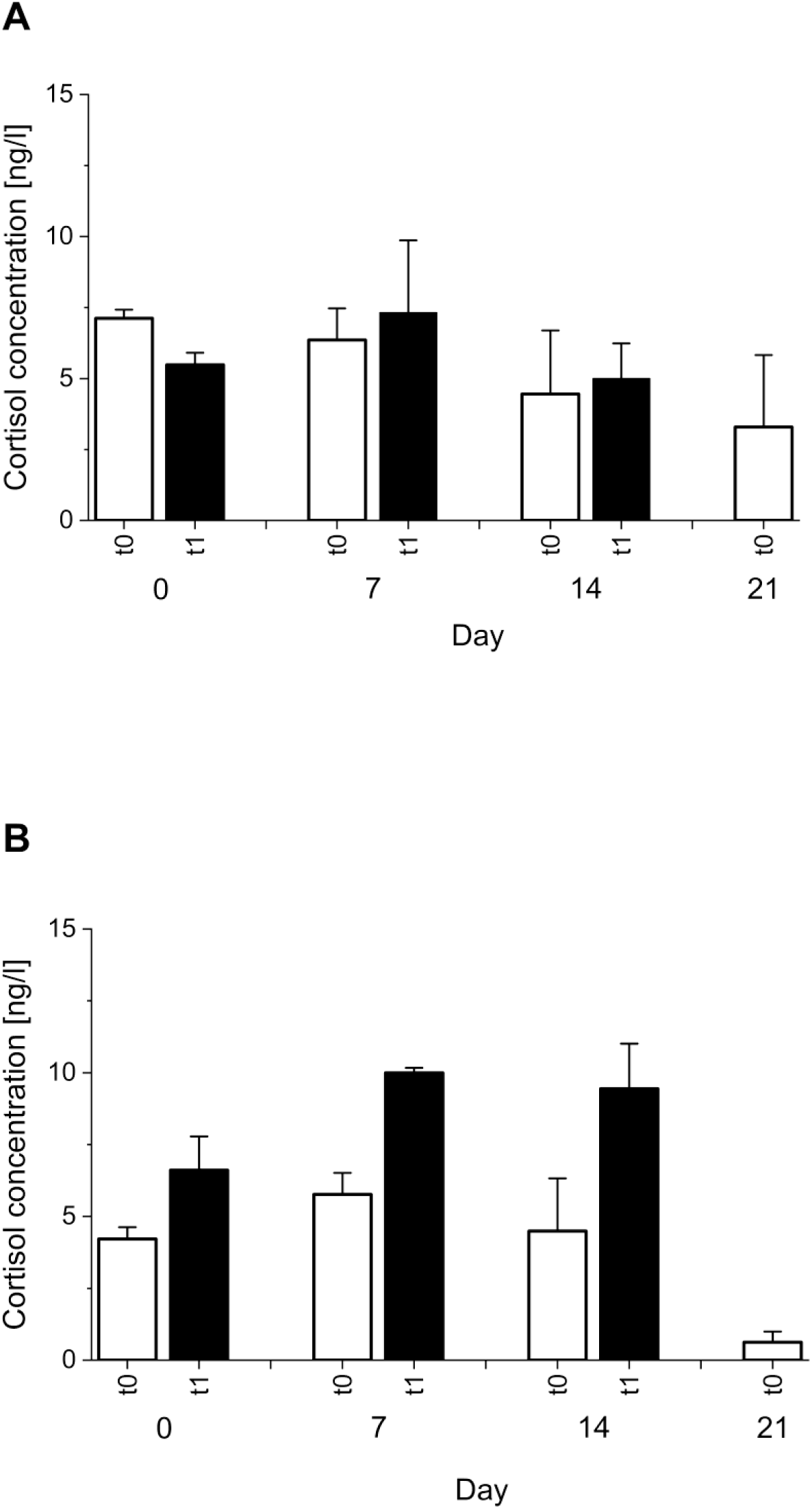
Averages and standard deviations for 2 technical replicates of the cortisol concentrations of the water samples from (A) the CAP setup and (B) the UV setup, taken at the outlets (CAP C and D, UV C and D) for the sampling times t_0_ (white columns) and t_1_ (black columns) of the respective sampling days

At the start of the trial (day 0), the initial cortisol concentration (before the CAP device was switched on (t_0_)) was 7.12 ng/l. Interestingly, the concentration decreased with the CAP treatment to 5.49 ng/l (t_1_). At the next sampling (day 7), the cortisol concentration only slightly, increased during treatment (from 6.36 ng/l to 7.32 ng/l, the difference was not significant). For the following sampling days, an overall decrease in the cortisol levels compared to the previous samplings was observed on day 14 (4.45 ng/l and 5.00 ng/l for t_0_ and t_1_) and on day 21 (3.29 ng/l for t_0_).

In contrast, the UV treatment, surprisingly led to an increase in cortisol. Although, the initial cortisol concentration (day 0, t_0_) was, with 4.22 ng/l, similar to the CAP setup, at the end of the day it had reached 6.61 ng/l for UV at t_0_ was in a similar range as in the CAP-treated setup (4.22 ng/l). For the time point t_1_ after 7 days the cortisol concentration before treatment slightly increased (up to 6.61 ng/l). Most pronounced was the difference at t_0_ and t_1_ of the UV setup day 7 and day 14, where cortisol levels almost doubled, which could be due to the circadian rhytm of cortisol or the increased stress level.

Thus, CAP did not increase the stress level of fish and further no behavioral changes were noticed in contrast to the UV setup. This indicates that CAP may serve as a beneficial tool to enhance fish welfare in aquacultures.

### 3.5 Energy efficiency

Energy requirements for both systems were derived from their respective wall-plug power and their time of operation during an experiment. The UV lamps with 55 W were switched on during the whole time of the experiment, corresponding to an energy of 2.64 kW for 24 h. The CAP device was operated in 10 min intervals with breaks of 30 min for altogether 8 hours. Accordingly, the duty time was 2.02 h in total each day. With a power of 500 W (± 20 W) this corresponds to a dissipated energy of 1.01 kW for the respective time of treatment per day. Despite the already 2.6 times lower energy requirement for the CAP treatment, it should still be mentioned that the unit is an experimental device without any optimization of energy efficiencies, whereas the UV treatment was conducted with a mature commercial system.

## 4. Discussion

A key issue for the cultivation of healthy fish in RAS is the control of microbial loads. Therefore, the decontamination of water in RAS is typically achieved with UV treatment or ozonation (Xiao et al. 2019). This state-of-the-art is apparently sufficient to meet current needs. However, these technologies have drawbacks. The inactivation via UV depends on the penetration depth of the light reaching the microorganisms (Zhu 2002) for the necessary direct interaction. In RAS the penetration can be limited due to turbidity caused by remaining suspended solids from feed or faeces. Aging of the lamps and fouling at the outer surface of the lamps further reduce the efficacy over time. The alternative use of ozonation is based on the oxidation process of cell membranes to ensure microbial inactivation, which is independent of water turbidity. However, ozone, as a strong oxidative agent can become toxic for the fish and also the useful nitrifying microorganisms in the biofilter (Schroeder et al. 2015). Even small residual concentrations have been found to damage gill tissues causing respiratory distress or even increase mortality in juvenile tilapia (*Oreochromis niloticus*) (Wuertz et al. 2023). Especially juvenile stages are rather sensitive in this respect. Moreover, the corrosive nature of ozone affecting system components, e.g., degrading plastics and rubber seals necessitate a regular replacement of spare parts. Accordingly, the use of ozone needs to be tightly controlled and carefully applied. Hence, ozonation is primarily applied in fresh-water aquacultures and, in general, not as widely used as UV-treatment (at least in Europe) (Stiller et al. 2020, Weidlich et al. 2026). More recently, UV treatments as well as ozonation have been indicated to increase antibiotic resistance genes (Xue et al. 2023), which points towards a growing concern for the spread of antibiotic resistances through aquacultures.

Therefore, novel approaches for disinfection in RAS, are still of interest to address the shortcomings of UV treatments and ozonation. CAP treatments offer such an alternative by providing short-lived reactive nitrogen species, such as peroxynitrite (Balazinski et al. 2021, Pogoda et al. 2026) , which are dissolving and efficient for the inactivation of microorganisms (independent from turbidity). However, plasma technologies need to guarantee fish welfare (or even improve it) and be economically competitive.

The most obvious advantage of the investigated CAP treatment, in comparison to the UV application, was, hence, the much lower energy requirement. For the CAP system a 2.5 times lower need of energy achieved a microbial load equal to the UV setup (CAP: 2.16 10^4^ cfu/ml; UV: 3.27·10^4^ cfu/ml). Reasonably, the shorter treatment time of in total 2.02 h for CAP in contrast to 24 h with UV provided the more energy-efficient method. Apparently, a more comprehensive, respectively faster microbial inactivation could be provided with CAP.

The determined microbiological results correspond to previously performed investigations under laboratory conditions, that also revealed comparable antimicrobial efficacies for both technologies (Balazinski et al. 2025). This applied especially to Gram-negative microorganisms, such as *Pseudomonas fluorescens* (with a logarithmic reduction factor (log_10_-RF) of 2.7 for CAP and 2.9 for UV) or *Aeromonas salmonicida* (with a log_10_-RF of 4.9 for CAP and 5.5 for UV, respectively). Altogether, the experiments conducted in an actual RAS, and previously in laboratory setups, show that consistent microbial loads can be achieved.

However, an unspecific inactivation of microorganisms is not necessarily beneficial to enhance fish welfare. Instead, a stable and healthy microbial community, i.e., microbiome, is essential for fish welfare e.g., protecting the fish from toxic residues, stimulating metabolic processes and promoting healing processes (Lorgen-Ritchie et al. 2023, Qi et al. 2026). While many studies assessed the microbiome of fish grown in aquacultures (gut, gills, skin etc.; (Berggren et al. 2022, El-Son et al. 2025, McMurtrie et al. 2025)), only a few studies considered changes of the water microbiome (Ruiz et al. 2025). Ruiz et al. investigated the microbiome of the mucosal organs of Rainbow trout and considered also the surrounding microbiome of the water. Many factors can affect the composition of the microbiome. Kanika et al. reviewed several impact factors for the fish gut microbiome, such as the fish taxa, feeding habits as well as habitat characteristics and geomorphology as influencing parameters for the respective phyla distribution of microorganisms (Kanika et al. 2025). For a closer look on the microbial community with respect to fish welfare, an in-depth analysis with the sequencing of the 16S rRNA amplicon was performed. The analysis distinguishes genes, which exist in all bacterial cells and consists of highly conserved and hypervariable regions. The variable regions refer specifically to individual bacterial species and enable the detection even in samples containing low biomass of bacteria (Jin et al. 2024). On the phyla level high abundances were found in our study for *Pseudomonadota* in both setups (CAP: 32%; UV: 57%). Further, the phyla *Bacteroidota* (CAP: 6% ± 3%; UV: 9% ± 4%) and *Acidobacteroidota* (CAP: 3% UV: 2%) were found, however to a far lower extent. The phyla *Pseudomonadota* and *Bacteroidota* were also found by Ruiz et al. with also high abundances (43%, 29%) in a RAS with cultivated Rainbow trout (the type of decontamination was not further specified). The high abundance of *Pseudomonadota* in our studies was also confirmed by the MALDI-TOF results. From the thirteen identified samples, nine were assigned to this phylum. The other samples were categorized as *Bacteroidota* and *Bacillota*. A rough agreement on the phyla distribution between our and the study of Ruiz et al. was found, except for *Bacillota*.

While the phyla level represents only a very general perspective, more information can be obtained from the detected genera. There are some genera which were mainly associated to diseases in fish, such as *Vibrio*, *Aeromonas* and *Shewanella* (Li et al. 2017). From these, *Vibrio* and *Shewanella* were not detected in any of our samples. *Aeromonas*, in contrast, was identified only in two UV samples on day 0 and 14, but only with low abundances (max. 3.2%), which did not further increase.

The weekly determination of the microbial community in the respective setups showed that the development was different. Thus, a broad variety of genera was found in both setups. Especially for CAP remarkable changes in the community were detected after one week. Considering the decontamination via UV in the adaption phase of the fish a change in the water microbiome was expected. The consistency of the community determined in the following sampling dates showed that the microbial community remained stable. Whereas, for the UV setup for which a constant microbial community was expected no such stable microbial community was noted within the trial as for CAP. Although, the same decontamination technology was applied in the adaption phase before more fluctuations have been detected in the distribution of the genera with higher abundances. Reasonable for this could be the fish development and therefore occurring changes in the water turbidity and also increasing amounts of excreted faeces and supplied fish feed. It would be conceivable that the degree of impact towards varying conditions is different between the technologies.

However, within the multitude of detected genera *Cetobacterium*, *Flavobacterium* and *Pseudomonas* were found in both setups (Tab. S II, Supplementary Information). *Cetobacterium* represents a genus, which plays a major role in fish metabolism. Bacteria of this genus are beneficial for fish health and are involved in processes, such as lipid and carbohydrate metabolization (Wang et al. 2025). The other two genera *Flavobacterium* and *Pseudomonas*, however, contain species which are associated with fish diseases, with *Flavobacterium columnare* causing the columnaris disease (Loch and Faisal 2015). Further, *Flavobacterium psychrophilum* and *Flavobacterium branchiophylum* were associated to cause fish diseases (cold water disease, bacterial gill disease) (Shotts and Starliper 2011, Starliper 2011). Within the genus *Pseudomonas*, *Pseudomonas fluorescens* represents the most important species causative for diseases on skin and fin (Pękala-Safińska 2018). The abundances of these genera underlaid fluctuations during the experiment, however, strongly decreasing in both setups during the trial. Thus, in this study the 16S rRNA gene amplicon sequencing was performed on 500 bp a detection was only possible for the genera. Therefore, it is not clear whether, if the mentioned pathogenic species were present or maybe non-pathogenic species dominated the genera that were found. Further microbiome studies, investigating more than 500 bp are necessary for species determination, which would allow for a comprehensive assessment especially for the genera *Flavobacterium* and *Pseudomonas*. Furthermore, the relatively high amount of hits with a low abundance below 2% (CAP: 47% ± 4%, UV: 42% ± 11%) shows, that the microbial community is even more complex than represented (Fig. S II, Supplementary Information) obtaining only higher abundances. However, still more research is recommended for a better understanding of microbial communities and their interlinking effect towards fish welfare and fish development to better classify detected changes.

Based on the sequencing of the 16S rRNA amplicon, changes to the microbial composition were further evaluated concerning α- and β -diversity. In general, the α-diversity describes the diversity within a community. The determination of this metric in this study allows to detect changes in the biodiversity, whether within the CAP or UV setup. The α-diversity, expressed by the Shannon-Index showed that in the respective setups no remarkable changes occurred within the experiment duration. However, the loss on the detected genera after one week (CAP: 25 to 12, UV: 23 to 15) indicated, that a treatment, whether by CAP or UV, affects the microbial community, which, after initial adaptation remained stable. The β-diversity whereas, is used to compare the biodiversity of different communities, here to compare the biodiversity in both setups (Cassol et al. 2025). The calculated β-diversity resulted in closely related values of both setups throughout the sampling dates, indicating that the microbial diversity was similar and no deviating effects occurred. Although, the α- and β-diversity confirmed a stable composition of the microbial community, especially the β-diversity is the indicator of main importance for the assessment of the impact that CAP has on the microbial community compared to UV. Nevertheless, an evaluation, if these changes are beneficial for the fish welfare or not is difficult. Some studies which investigated the microbiome diversity in aquaculture plants indicated, that a diverse microbiome was associated with improved fish health and welfare (Infante-Villamil et al. 2019). Differences regarding the diversity were found, when comparing the water microbiome with the host-associated microbiome of vertebrates and invertebrates (Infante-Villamil et al. 2019). While a high microbiome diversity was found in the water, this did not reflect the host-associated microbiome. The water microbiome is, in principle, also affected by stress levels of the animals, since numbers of beneficial microorganisms decrease with rising stress values (Boutin et al. 2013). However, no related changes to the microbiome could be determined either for UV or CAP treatments. Therefore, cortisol concentrations were measured from water samples, to determine, if the decontamination devices affected the fish at a hormonal level, i.e., impose stress. Although, cortisol measurements from water samples provide lower concentrations than the usual measurements carried out with blood plasma or sampling of scales and fins it was shown that after a time-delay the changes can also be detected in water samples (Ellis et al. 2004, Gabor and Contreras 2012, Knopf et al. 2014). While cortisol concentrations were slightly lower for CAP treatments (CAP: 5.6 ng/l; UV: 5.9 ng/l), the differences between the setups and also the comparison of the values prior to CAP treatment and after starting the treatment were not significant, indicating that CAP does not promote a stress response. Ideally, the intermittent operation of the CAP unit may be gentler than the continuous application of UV. Accordingly, no differences were observed in the gill movement, the swim behavior and overall behavior of the fish in both setups. Therefore, a score of 0 applied to the respective categories of the scoresheet (see Tab. SI, Supplementary Information), wherefore no intervention was necessary.

Concurrently, values for pH, temperature, conductivity, and oxygen were consistent in both setups (except for the technical failure of the data logger in the first week of the UV setup). While the pH value was relatively high at the beginning (almost 9), this value decreased during the experimental trial in both setups. This decrease is common in RAS with rising stocking densities due to the increased biological respiration and decomposition of organic matter leading to higher oxygen consumption. This further promotes the release of carbon dioxide, which in the following results in lower pH values (Li et al. 2023). Although it is known, that the species generated by CAP can also provide a decrease of the pH value and a rise of the conductivity, no striking differences in comparison to the UV setup were determined. These results contrast previously conducted laboratory experiments where the treatment of a much smaller water volume of 90 l did give rise to higher pH values and higher conductivities (Balazinski et al. 2025). Conversely, for the treatment of a volume of 680 l in the RAS, reactive species that are produced by the CAP treatment (and responsible for these changes), are apparently better distributed (and diluted). While, the measured oxygen levels, with a mean of approximately 10 mg/l is referred to normoxic (Policar et al. 2024) the temperature indifference, also previously observed in small-scaled experiments (performed with 90 l) offered stable conditions. Furthermore, water in aquacultures contains organic matter, such as fish faeces or feed residues, which could promote scavenging reactions for reactive species, which was previously shown for plasma-treated liquids (Hahn et al. 2019, Lavrikova et al. 2025).

In conclusion, the concern raised by laboratory experiments that the production of reactive species, while responsible for the intended disinfection, could negatively affect the microbiome and fish welfare was not shown to be relevant in the actual RAS. Concentrations of nitrate and nitrate, which are known to be generated by the CAP device in significant amounts, were quantified by ion chromatography. The analysis showed that there is a noticeable increase, especially for nitrite when the CAP treatment was started. Nitrite can become harmful to fish, affecting the mechanism of chloride/hydrogen carbonate ion exchange, which in the following can reduce the oxygen-carrying capacity of blood causing, e.g., tissue hypoxia metabolism (Jie et al. 2026). The Rainbow trout, cultivated in RAS, is a strain from the local longstanding breeding program (Borchel 2015) and adapted to brackish water conditions, hence, changes in water conditions (Rebl et al. 2013, Verleih et al. 2015). However, salmonid species, such as Rainbow trout, are known to be susceptible to increasing nitrite values (Jensen 2003). To minimize the production of nitrate and nitrite, without compromising disinfection, the treatment duration was adjusted during the first day of the experimental trial, to the interval of 10 min treatment, 30 min pause. This intermittent operation allowed to generate sufficient amounts of reactive species for an effective decontamination, likewise leaving enough time for the microorganisms (especially the nitrite oxidizing ones) in the biofilter to oxidize nitrite to nitrate. This balanced treatment pattern ensured stable nitrite levels, which did not harm the fish in our experiments.

Altogether, the results confirm CAP as a competitive alternative to UV or ozonation for the disinfection in RAS. Microbial loads can be maintained without relevant changes to the microbiome or adverse effects on fish welfare. The technology further offers some conceptional advantages. Since the inactivation is based on the introduction of short-lived radical species in the water, it is not prone to insufficiencies of a required direct interaction of UV light, which can be made difficult in turbid water or deposits on lamp housings. With respect to ozonation, it avoids the needed, often cumbersome, infrastructure and difficulties for production, delivery and especially dosing. Instead, the electrical operation of CAP in ambient air permits a control that can be adjusted to actual needs, depending on microbial loads and water parameters that are monitored anyway. This was already exploited here to establish an intermittent operation with the additional benefit of reduced operational costs. Concurrently, the method is inherently scalable. Two pipes were used in parallel for the treatment in the investigation but several more could be added depending on required treatment volumes. (This concept of parallelization is likewise pursuit for larger scale ozone generators.) If a centralized treatment of the entire water in a RAS or an installation of smaller units for individual tanks (similar to the UV treatment) is the better approach, still needs to be evaluated. This will also guide the further technological development, which should include the so far neglected optimization of energy efficiencies.

The exploitation of explicitly transient reactive species, which is an advantage with respect to fish welfare, is also an environmental benefit since no harmful potentially toxic substances are released, for example, with waste waters. (With a working biofilter not even relevant amounts of nitrate and nitrite.) This extends to the built of the systems. Unlike for the currently used UV lamps, which need mercury for operation, no problematic materials are needed for CAP devices.

## 5. Conclusion

The presented pilot study demonstrated the potential of plasma treatment for the decontamination of water in recirculating aquaculture systems. The same hygienic standard was achieved as for a standard UV treatment and no adverse effects on fish welfare have been determined. However, the scope of the trial was limited to Rainbow trout as one particular species of fish. Also, treatments were carried out for a short period of 3 weeks with even shorter daily operation of only 2 hours. Despite and regardless of the remarkable results for this design, more praxis-relevant investigations need to be conducted to gain further knowledge on the processes, its benefits, but also challenges for an introduction into markets. Especially long-term experience is necessary, for example, with respect to process stability and maintenance requirements. The possibilities for the control of the treatment, e.g., duty cycles of operation, might be challenging but also provide an opportunity to address variations between different aquaculture settings, determine requirements and impact on other fish species, and also for larger or smaller volumes of water and stocking densities.

## Acknowledgements

Rica Tielebier, Andreas Tielebier, Daniel Genz and Stefan Herper is thanked for technical assistance during animal experiments. Dominik Schneider is thanked for providing information on sample preparation for amplicon sequencing. The plasma sources department of the Leibniz institute for Plasma Science and Technology e.V. is thanked for the construction of the plasma source.

## Supplementary Information

**Tab. S I.** Scoresheet for the assessment of the fish welfare including instructions for action.

| **Category** | **Degree** | **Score** | **Instruction** |
|---|---|---|---|
| Gill movement during/after treatment | Normal | 0 |  |
|  | Slightly shallowed | 1 | Manual movement of the fish in the tank. Targeted monitoring at the next checkup |
|  | Considerably shallowed | 2 | Immediate „resuscitation“, flushing the gills with fresh tank water by a hose |
|  | Considerably shallowed for longer than 10 min | 3 | Termination, if ventilatory measures do not result in improvement after 10 min |
| Swim behavior during/ after treatment | Normal | 0 |  |
|  | Staggering | 1 | Continued observation of the fish |
|  | Lateral position | 2 | Continued observation of the fish |
|  | Supine position | 3 | Termination |
| Overall changes in behavior during/ after treatment | None | 0 |  |
|  | Restlessness | 1 | Continued observation of the fish |
|  | Loss of postural reflex | 3 | Termination |
|  | Panicked, fast swimming for no reason | 3 | Termination |

**Tab. S II.** Summation of the relative abundances of the identified phyla, family and genera of the respective sampling positions for the CAP and the UV setup of the sampling days (A= day 0, B= day 7, C= day 14, D= day 21).

| **CAP** |  |  |  | **UV** |  |  |  |
|---|---|---|---|---|---|---|---|
| Phylum | Family | Genus | Relative abundance* [%] | Phylum | Family | Genus | Relative abundance* [%] |
| *Acidobacteriota* | *Holophagaceae* | *Incertae* | 1.02804 | *Acidobacteriota* | *Holophagaceae* | *Incertae* | 1.66271 |
|  | *Vicinamibacteraceae* | *Incertae* | 2.85861 | *Actinomycetota* | *Nocardiaceae* | *Rhodococcus* | 10.93011 |
| *Bacteroidota* | *Flavobacteriaceae* | *Flavobacterium* | 1.58706 |  | *Nocardiaceae* | *Nocardia* | 3.27341 |
| *Chloroflexota* | *Incertae* | */* | 15.37101 | *Bacillota* | *Clostridiaceae* | *Clostridium* | 7.60095 |
|  | *JG30-KF-CM45* | *Incertae* | 2.75190 | *Bacteroidota* | *Flavobacteriaceae* | *Flavobacterium* | 2.56519 |
| *Deinococcota* | *Deinococcaceae* | *Deinococcus* | 1.26965 |  | *Spirosomataceae* | *Emticicia* | 0.98645 |
| *Fusobacteriota* | *Fusobacteriaceae* | *Cetobacterium* | 5.57901 | *Chloroflexota* | *Incertae* | */* | 5.28267 |
| *Nitrospirota* | *Nitrospiraceae* | *Nitrospira* | 11.85099 |  | *JG30-KF-CM45* | *Incertae* | 1.92663 |
| *Pseudomonadota* | *Alteromonadaceae* | *Rheinheimera* | 6.08115 | *Fusobacteriota* | *Fusobacteriaceae* | *Cetobacterium* | 7.81127 |
|  | *Azospirillaceae* | *Azospirillum* | 0.98131 | *Nitrospirota* | *Nitrospiraceae* | *Nitrospira* | 1.16313 |
|  | *Cellvibrionaceae* | *Cellvibrio* | 2.41609 | *Pseudomonadota* | *Aeromonadaceae* | *Aeromonas* | 3.24624 |
|  | *Caulobacteraceae* | *Brevundimonas* | 1.41745 |  | *Alteromonadaceae* | *Rheinheimera* | 7.64660 |
|  | *Comamonadaceae* | *Acidovorax* | 2.05440 |  | *Azospirillaceae* | *Azospirillum* | 1.07479 |
|  | *Comamonadaceae* | *Hydrogenophaga* | 3.91643 |  | *Cellvibrionaceae* | *Cellvibrio* | 1.28092 |
|  | *Comamonadaceae* | *unclassified* | 3.18287 |  | *Caulobacteraceae* | *Caulobacter* | 2.91519 |
|  | *Comamonadaceae* | *Variovorax* | 6.85091 |  | *Comamonadaceae* | *Acidovorax* | 1.62901 |
|  | *Hyphomicrobiaceae* | *Hyphomicrobium* | 2.14635 |  | *Comamonadaceae* | *unclassified* | 5.43679 |
|  | *Lysobacteraceae* | *Arenimonas* | 3.11170 |  | *Lysobacteraceae* | *Arenimonas* | 1.82106 |
|  | *Oxalobacteraceae* | *Undibacterium* | 0.91146 |  | *Paracoccaceae* | *Fuscovulum* | 1.68910 |
|  | *Paracoccaceae* | *Fuscovulum* | 4.62068 |  | *Pseudomonadaceae* | *Pseudomonas* | 21.04687 |
|  | *Paracoccaceae* | *unclassified* | 1.54172 |  | *Rhodocyclaceae* | *Dechloromonas* | 2.58643 |
|  | *Pseudomonadaceae* | *Pseudomonas* | 10.0076 |  | *Sphingomonadaceae* | *Sphingobium* | 1.43633 |
|  | *Reyranellaceae* | *Reyranella* | 1.99377 |  | *Yersiniaceae* | *unclassified* | 2.77118 |
|  | *Sphingomonadaceae* | *Sphingobium* | 2.86815 | *Thermodesulfobacteriota* | *Desulfovibrionaceae* | *Fundidesulfovibrio* | 2.21694 |
|  | *Xanthobacteraceae* | *Xanthobacter* | 1.16822 |  |  |  |  |
| *Verrucomicrobiota* | *Rubritaleaceae* | *Luteolibacter* | 2.43349 |  |  |  |  |
| **CAP** |  |  |  | **UV** |  |  |  |
| Phylum | Family | Genus | Relative abundance* [%] | Phylum | Family | Genus | Relative abundance* [%] |
| *Acidobacteriota* | *Vicinamibacteraceae* | *Incertae* | 2.07310 | *Actinomycetota* | *Nocardiaceae* | *Rhodococcus* | 1.37367 |
| *Bacteroidota* | *Flavobacteriaceae* | *Flavobacterium* | 4.28683 |  | *Nocardiaceae* | *Nocardia* | 3.59033 |
|  | *Saprospiraceae* | *Incertae* | 4.41893 | *Bacteroidota* | *Flavobacteriaceae* | *Flavobacterium* | 3.15036 |
| *Chloroflexota* | *Incertae* | */* | 28.83075 |  | *Spirosomataceae* | *Arcicella* | 3.39071 |
|  | *JG30-KF-CM45* | *Incertae* | 0.96209 |  | *Spirosomataceae* | *Emticicia* | 3.41771 |
| *Fusobacteriota* | *Fusobacteriaceae* | *Cetobacterium* | 2.09892 | *Chloroflexota* | *Incertae* | */* | 9.17023 |
| *Nitrospirota* | *Nitrospiraceae* | *Nitrospira* | 13.85711 | *Fusobacteriota* | *Fusobacteriaceae* | *Cetobacterium* | 15.93319 |
| *Patescibacteria* | *Incertae* | */* | 2.83339 | *Pseudomonadota* | *Alteromonadaceae* | *Rheinheimera* | 1.07807 |
| *Pseudomonadota* | *Comamonadaceae* | *unclassified* | 1.17288 |  | *Azospirillaceae* | *Azospirillum* | 2.14789 |
|  | *Hyphomicrobiaceae* | *Hyphomicrobium* | 2.35183 |  | *Caulobacteraceae* | *Caulobacter* | 1.95114 |
|  | *Paracoccaceae* | *Fuscovulum* | 10.16144 |  | *Comamonadaceae* | *Acidovorax* | 9.20319 |
|  | *Paracoccaceae* | *unclassified* | 2.21504 |  | *Comamonadaceae* | *unclassified* | 16.11141 |
|  | *Pseudomonadaceae* | *Pseudomonas* | 7.08966 |  | *Comamonadaceae* | *Limnohabitans* | 1.96754 |
| *Verrucomicrobiota* | *Rubritaleaceae* | *Luteolibacter* | 13.45876 |  | *Pseudomonadaceae* | *Pseudomonas* | 18.65650 |
| *Unclassified* | *Unclassified* | *Unclassified* | 4.18928 |  | *Sphingomonadaceae* | *Sphingobium* | 4.94782 |
|  |  |  |  | *Verrucomicrobiota* | *Rubritaleaceae* | *Luteolibacter* | 3.91022 |
| **CAP** |  |  |  | **UV** |  |  |  |
| Phylum | Family | Genus | Relative abundance* [%] | Phylum | Family | Genus | Relative abundance* [%] |
| *Acidobacteriota* | *Pyrinomonadaceae* | *Incertae* | 2.56080 | *Actinomycetota* | *Nocardiaceae* | *Rhodococcus* | 1.71181 |
|  | *Vicinamibacteraceae* | *Incertae* | 0.89025 |  | *Nocardiaceae* | *Nocardia* | 11.22943 |
| *Bacteroidota* | *Flavobacteriaceae* | *Flavobacterium* | 4.50811 | *Bacillota* | *Clostridiaceae* | *Clostridium* | 2.06518 |
|  | *Saprospiraceae* | *Incertae* | 1.18214 | *Bacteroidota* | *Flavobacteriaceae* | *Flavobacterium* | 11.30807 |
| *Chloroflexota* | *Incertae* | */* | 32.55915 |  | *Spirosomataceae* | *Lacihabitans* | 1.29797 |
| *Fusobacteriota* | *Fusobacteriaceae* | *Cetobacterium* | 2.47367 | *Chloroflexota* | *Incertae* | */* | 3.48266 |
| *Nitrospirota* | *Nitrospiraceae* | *Nitrospira* | 6.38174 | *Fusobacteriota* | *Fusobacteriaceae* | *Cetobacterium* | 4.29696 |
| *Patescibacteria* | *Incertae* | */* | 3.38509 | *Plantomycetota* | *Pirellulaceae* | *Pir4* | 2.35560 |
| *Pseudomonadota* | *Comamonadaceae* | *unclassified* | 2.68389 | *Pseudomonadota* | *Aeromonadaceae* | *Aeromonas* | 2.51694 |
|  | *Hyphomicrobiaceae* | *Hyphomicrobium* | 3.13741 |  | *Azospirillaceae* | *Azospirillum* | 1.14396 |
|  | *Paracoccaceae* | *Fuscovulum* | 8.57328 |  | *Caulobacteraceae* | *Caulobacter* | 2.58712 |
|  | *Paracoccaceae* | *unclassified* | 5.72882 |  | *Comamonadaceae* | *Acidovorax* | 2.69011 |
|  | *Pseudomonadaceae* | *Pseudomonas* | 6.63530 |  | *Comamonadaceae* | *Limnohabitans* | 6.86378 |
| *Verrucomicrobiota* | *Rubritaleaceae* | *Luteolibacter* | 16.81932 |  | *Comamonadaceae* | *unclassified* | 17.50372 |
| *Unclassified* | *Unclassified* | *Unclassified* | 2.48103 |  | *Comamonadaceae* | *Variovorax* | 1.18510 |
|  |  |  |  |  | *Enterobacteriaceae* | *Escherichia-Shigella* | 5.29743 |
|  |  |  |  |  | *Lysobacteraceae* | *Arenimonas* | 3.22685 |
|  |  |  |  |  | *Paracoccaceae* | *unclassified* | 6.20243 |
|  |  |  |  |  | *Paracoccaceae* | *Tabrizicola* | 2.03291 |
|  |  |  |  |  | *Pseudomonadaceae* | *Pseudomonas* | 7.48483 |
|  |  |  |  |  | *Sphingomonadaceae* | *Sphingobium* | 1.54875 |
|  |  |  |  | *Verrucomicrobiota* | *Pedosphaeraceae* | *Incertae* | 1.96838 |
| **CAP** |  |  |  | **UV** |  |  |  |
| Phylum | Family | Genus | Relative abundance* [%] | Phylum | Family | Genus | Relative abundance* [%] |
| *Acidobacteriota* | *Pyrinomonadaceae* | *Incertae* | 4.34634 | *Actinomycetota* | *Nocardiaceae* | *Rhodococcus* | 2.52441 |
| *Bacteroidota* | *Flavobacteriaceae* | *Flavobacterium* | 3.25199 |  | *Nocardiaceae* | *Nocardia* | 12.29661 |
|  | *Saprospiraceae* | *Incertae* | 5.03124 | *Bacillota* | *Staphylococcaceae* | *Macrococcus* | 1.41883 |
| *Chloroflexota* | *Incertae* | */* | 28.89946 | *Bacteroidota* | *Flavobacteriaceae* | *Flavobacterium* | 8.80667 |
| *Fusobacteriota* | *Fusobacteriaceae* | *Cetobacterium* | 2.03804 | *Chloroflexota* | *Incertae* | */* | 4.39959 |
| *Nitrospirota* | *Nitrospiraceae* | *Nitrospira* | 7.27471 | *Fusobacteriota* | *Fusobacteriaceae* | *Cetobacterium* | 9.25063 |
| *Patescibacteria* | *Incertae* | */* | 4.90486 | *Plantomycetota* | *Pirellulaceae* | *Pir4* | 2.02232 |
| *Pseudomonadota* | *Comamonadaceae* | *unclassified* | 2.74785 | *Pseudomonadota* | *Alteromonadaceae* | *Rheinheimera* | 1.36398 |
|  | *Hyphomicrobiaceae* | *Hyphomicrobium* | 2.22396 |  | *Burkholderiaceae* | *Limnobacter* | 1.90609 |
|  | *Paracoccaceae* | *Fuscovulum* | 10.38819 |  | *Caulobacteraceae* | *Caulobacter* | 1.58819 |
|  | *Paracoccaceae* | *unclassified* | 6.37824 |  | *Comamonadaceae* | *Aquabacterium* | 2.67190 |
|  | *Pseudomonadaceae* | *Pseudomonas* | 1.31661 |  | *Comamonadaceae* | *Limnohabitans* | 11.52715 |
| *Verrucomicrobiota* | *Rubritaleaceae* | *Luteolibacter* | 15.13726 |  | *Comamonadaceae* | *unclassified* | 15.50420 |
| *Unclassified* | *Unclassified* | *Unclassified* | 6.06123 |  | *Elsteraceae* | *Elstera* | 1.81241 |
|  |  |  |  |  | *Lysobacteraceae* | *Arenimonas* | 4.30033 |
|  |  |  |  |  | *Paracoccaceae* | *unclassified* | 2.55859 |
|  |  |  |  |  | *Paracoccaceae* | *Fuscovulum* | 2.11530 |
|  |  |  |  |  | *Paracoccaceae* | *Tabrizicola* | 1.95258 |
|  |  |  |  |  | *Pseudomonadaceae* | *Pseudomonas* | 9.65574 |
|  |  |  |  | *Verrucomicrobiota* | *Rubritaleaceae* | *Luteolibacter* | 2.32450 |

**Fig. S I.**
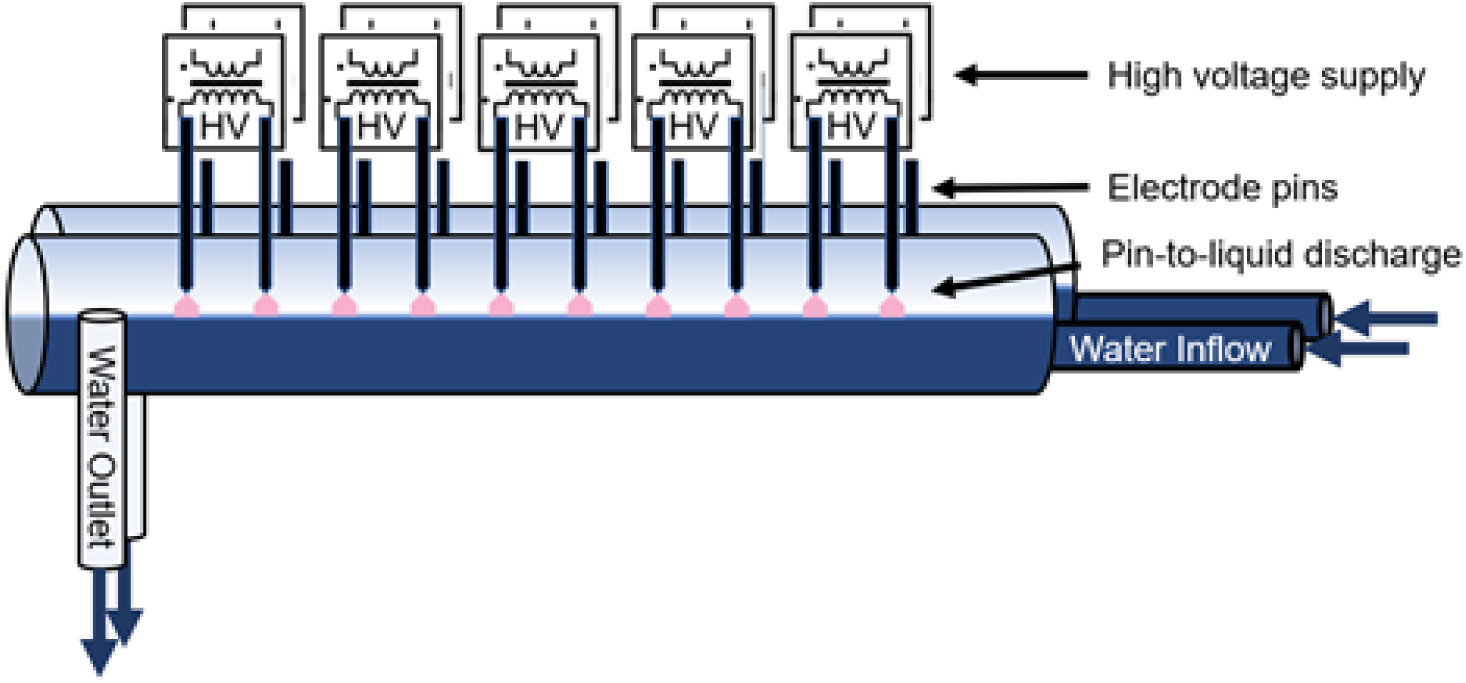
Schematic overview of the CAP decontamination unit.

**Fig. S II.**
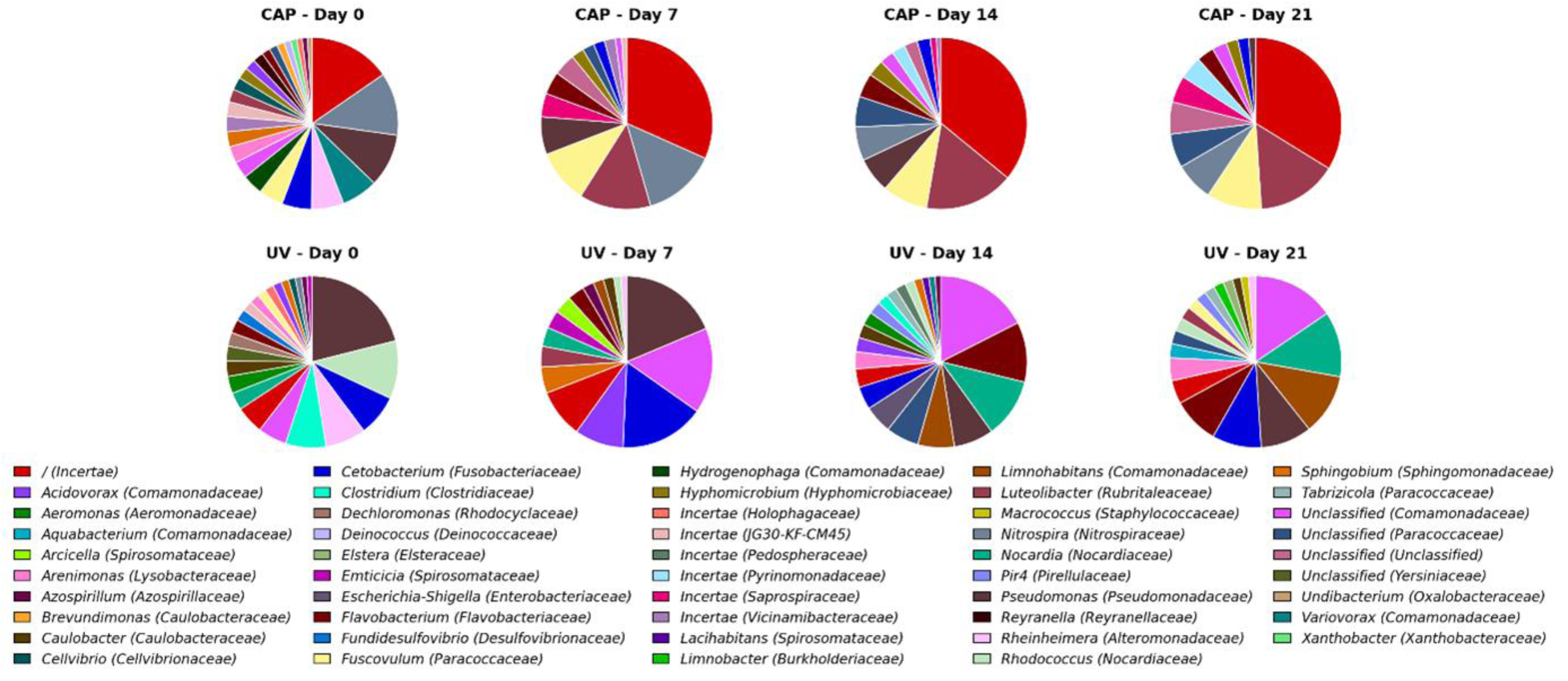
Relative abundances of the detected genera (with family) of the stratified data for CAP (top) and UV (bottom) of the respective sampling days (see 2.1.3).

**Fig. S III.**
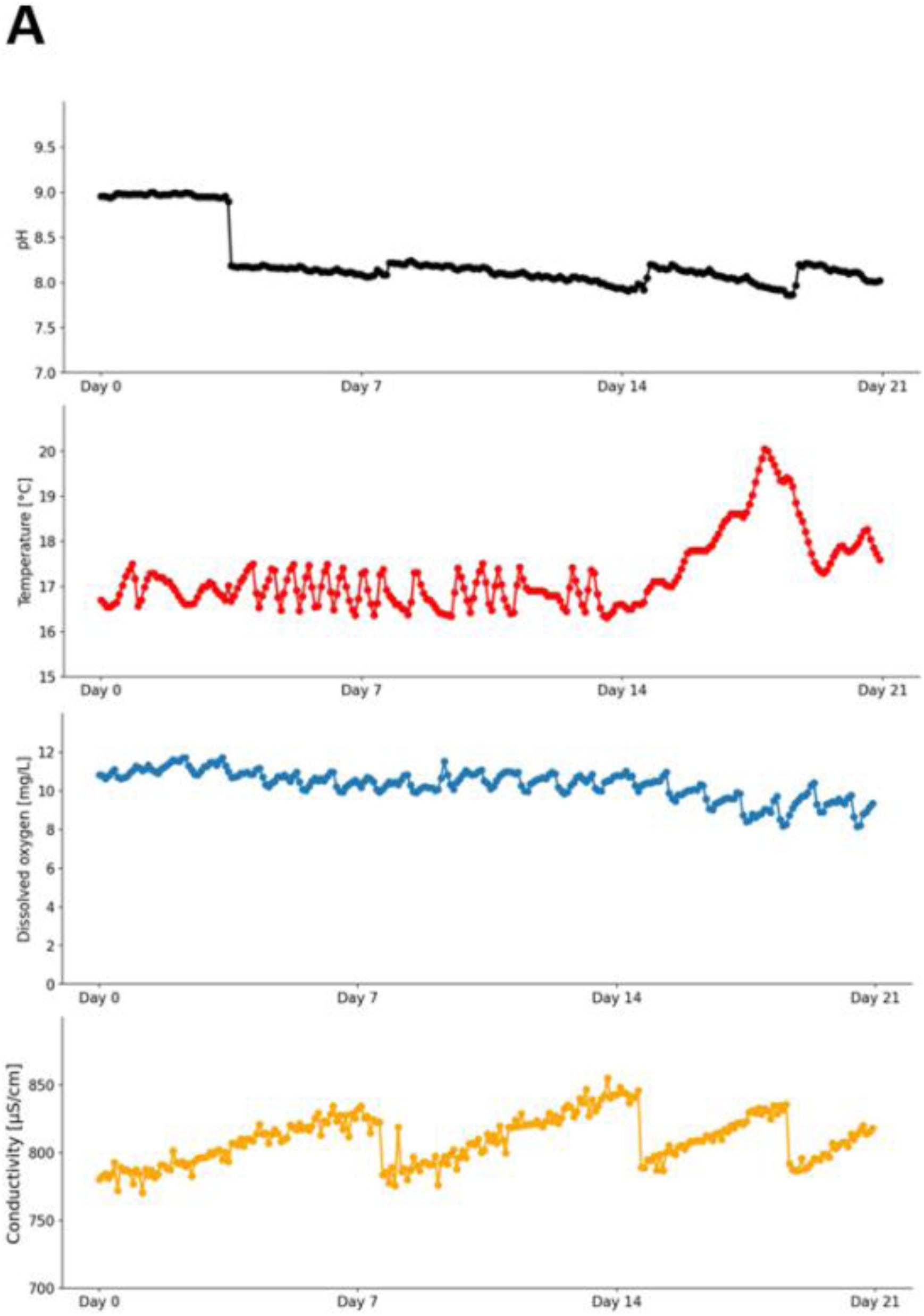
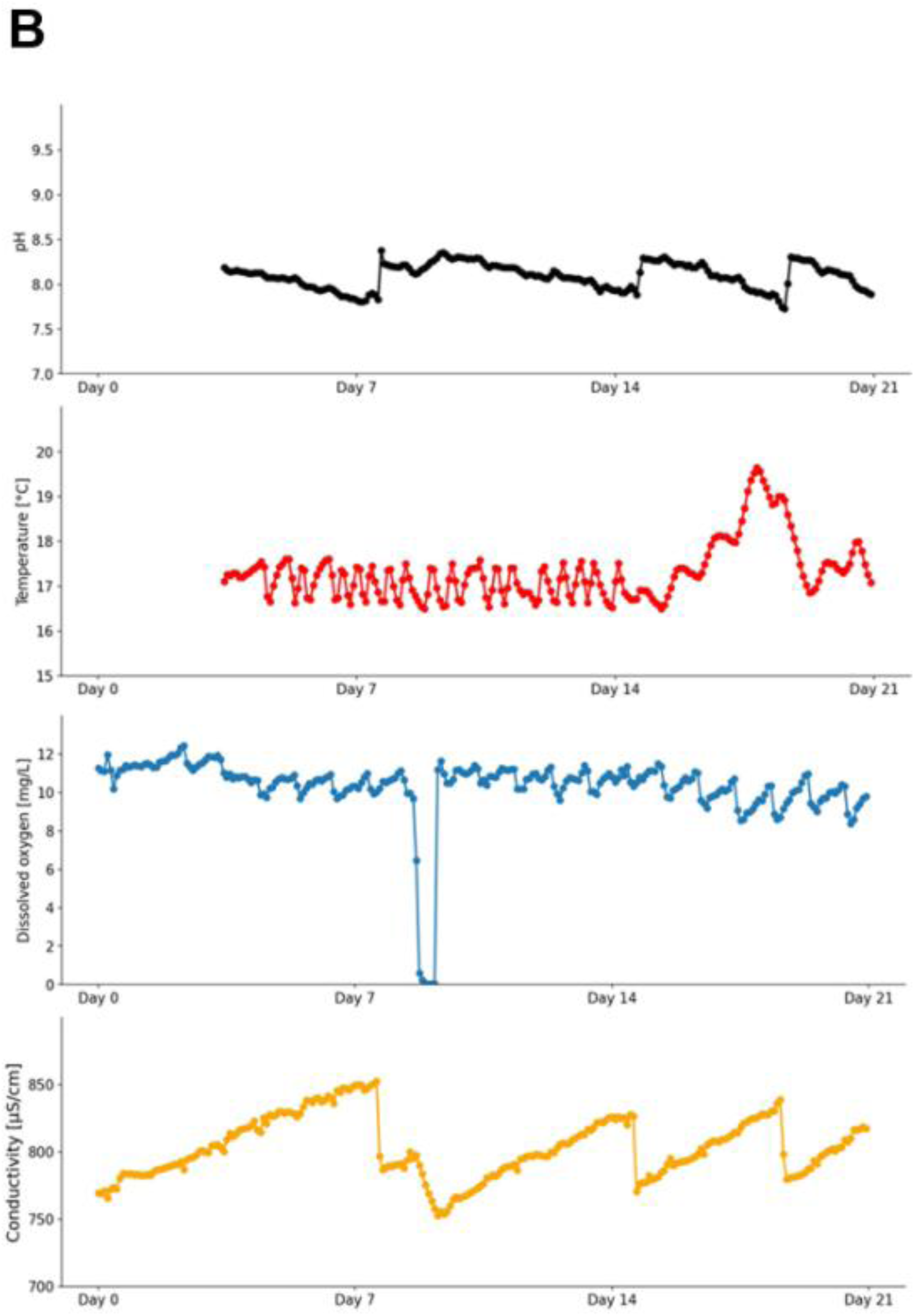
Values of pH, temperature, dissolved oxygen and conductivity for (A) the CAP setup and (B) the UV setup for the respective sampling days.

